# Loop stacking organizes genome folding from TADs to chromosomes

**DOI:** 10.1101/2022.07.13.499982

**Authors:** Antonina Hafner, Minhee Park, Scott E. Berger, Elphège P. Nora, Alistair N. Boettiger

## Abstract

While population level analyses reveal significant roles for CTCF and cohesin in mammalian genome organization, their contribution to chromatin structure and gene regulation at the single-cell level remain incompletely understood ^1–4^. Here, we use a super-resolution microscopy approach, Optical Reconstruction of Chromatin Architecture (ORCA) ^5^ to measure the effects of removal of CTCF or cohesin on genome folding across genomic scales. In untreated embryonic stem cells, we observe intricate, frequently stacked loops of chromatin which are largely dissolved upon cohesin removal. The loops compact chromatin at the < 3 Mb scale, increasing proximity between sequences not only within but also between TADs. We find multi-way contacts among loop anchors, preferentially at TAD borders, and these hubs largely dissolve upon CTCF degradation. CTCF-hubs bridge intervening TAD boundaries while keeping border distal regions from neighboring TADs apart outside the hub. Cohesin dependent loops at the < 3 Mb scale impede mixing at larger chromosomal scales through steric effects of loop stacking, dramatically reducing genomic cross-talk. Disruption of this ordered chromosomal structure led to increased cell-cell variability in gene expression, exceeding changes to average expression. Together our data revise the TAD-centric understanding of CTCF and cohesin, and provide a multi-scale, structural picture of how they organize the genome on the single-cell level through distinct contributions to loop stacking.

## Introduction

Mammalian genomes are organized in 3D into Topologically Associating Domains (TADs). TADs were discovered by Hi-C ^6–8^ and are defined as regions in which within-domain contacts are substantially more common than between-domain contacts ^9–12^. These preferential interactions within TADs are thought to be important for the specificity of gene regulation as enhancers and their target promoters frequently lie within the same TAD, and in a multitude of cases mutations disrupting TAD boundaries alter normal enhancer-promoter contact and gene expression ^1,3,13–15^.

Cohesin and CTCF are essential for the establishment and maintenance of TADs. Indeed, CTCF and cohesin binding is enriched at TAD boundaries and acute depletion of these proteins has been shown by Hi-C and imaging to lead to loss of TADs ^16–20^. Polymer modeling work has proposed that these two proteins work together in forming TADs through distinct roles; cohesin extrudes loops of chromatin but stalls at sites bound by CTCF ^21,22^, and considerable evidence supports these molecular roles ^2^. However, a detailed view of the physical 3D structure and how it is perturbed upon removal of CTCF or cohesin, with resolution from genes to whole chromosomes, is lacking. Such a high-resolution view may tease apart mechanisms or regimes in which CTCF and cohesin operate that are not distinguished by ensemble averages alone, and thus help us better understand how they effect cis-regulation of transcription. Microscopy approaches are well suited for investigating heterogeneity at the single-cell level.

Our earlier work on cohesin depleted cells imaging megabase-scale regions found ‘TAD-like’ domains were preserved in the absence of cohesin, though the position of their borders became more uniform ^20^. Two more recent microscopy studies that labeled TADs and looked at the consequences of cohesin and CTCF loss between and within TADs came to opposing conclusions ^23,24^. Upon degrading cohesin, Luppino and colleagues found reduced overlap between adjacently labeled TADs, while the volume of the TADs remained the same. By contrast, the second imaging study by Szabo and colleagues found that loss of cohesin did not significantly change the overlap between adjacently labeled TADs but increased the volume of each TAD.

To better understand the role of cohesin and CTCF on genome structure, we imaged genome structure from the few-kilobase scale of cis-regulatory elements to the scale of whole chromosomes, using a multi-scale chromosome tracing approach, similar to (refs ^25–27^). We multiplexed this imaging by using single-cell barcodes to combine different genetically edited cell lines and different treatment conditions into a single experiment, enabling direct measure of 3D structure in nanometers without risk of batch effects. Our single-chromosome data provide a detailed view of how cohesin and CTCF shape the 3D structure of chromatin from the kilobase to the whole chromosome scale; enhancing local contact by forming heterogeneous loops, preventing distal contact by loop clash, and facilitating border bypass by positioning loop anchors.

## Results

### We use microscopy to understand how TADs are lost upon cohesin and CTCF depletion

We first wanted to find out *how* TADs disappear; is it through loss of preferential interactions within TADs or gain of interactions between TADs? (**Fig. 1a**). These two scenarios would lead to distinct biophysical structures (**Fig. 1a**). We selected two ∼2.5 Mb regions, encompassing multiple TADs and sub-TAD structures (**Fig. 1d**), and containing developmentally important transcription factors, *Hoxa or Sox2*.

**Figure 1.**
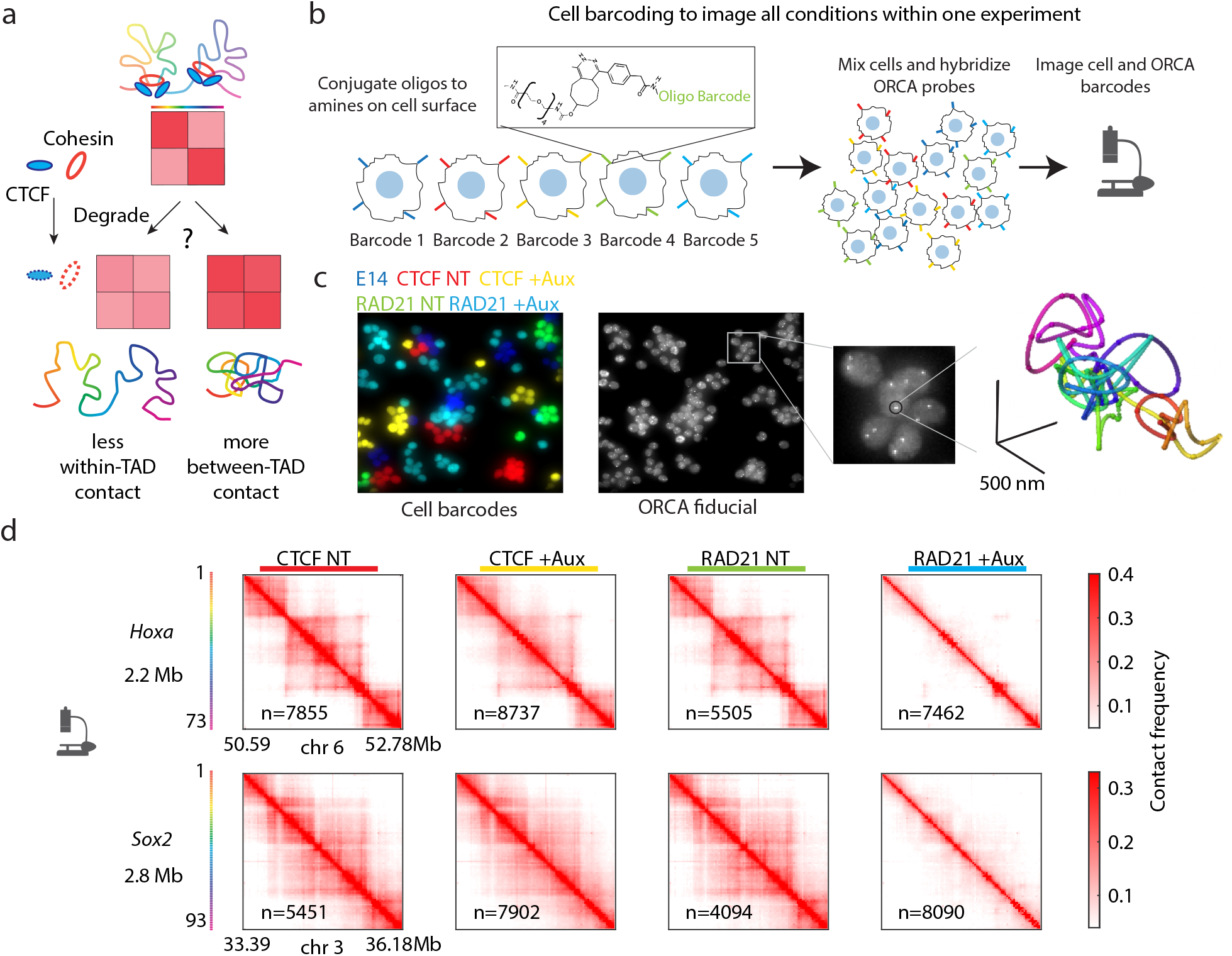
A multiplexed approach for chromosome tracing. **a.** Schematic depiction of two ways that preferential contacts within TADs can be lost upon degradation of CTCF or cohesin. **b**. Schematic depiction of the multiplexed ORCA workflow. **c**. Example of a field of view with pseudo-colored cell barcode labels for all imaged conditions: mES parental E14 cell line, CTCF-AID and RAD21-AID auxin-treated (+Aux) or untreated (NT) cells. Fiducial signal (labeling all probes in the Sox2 2.8 Mb domain), is shown as well as an example trace of a single chromosome for the Sox2 domain. **d**. Contact frequency maps from the demultiplexed cell conditions for both the *Hoxa* (2.2 Mb) and *Sox2* (2.8 Mb) regions imaged in 30 kb steps resulting in 73 steps for *Hoxa* and 93 steps for *Sox2*.

We used a microscopy chromosome tracing approach, Optical Reconstruction of Chromatin Architecture (ORCA) ^5^ to image single chromosomes in mouse embryonic stem cells (mESCs) with or without CTCF or RAD21, a subunit of the cohesin complex. We used cell lines where RAD21 and CTCF are tagged with the auxin inducible degron (AID) at their endogenous loci and can be rapidly depleted by addition of auxin (**Extended Data Fig. 1a**) ^19,24^. For all experiments described hereinafter, auxin-treated cells (+Aux) were treated with auxin for 4 hours, and control cells were not treated (NT).

To image all conditions (CTCF NT, CTCF +Aux, RAD21 NT, RAD21 +Aux) within the same experiment, we developed a multiplexed ORCA approach. We adapted a recently published method from single-cell sequencing ^28^ to attach distinct oligonucleotide-barcodes to the cell surface, after which cells can be combined, hybridized with ORCA probes, and imaged within the same experiment, where cell surface barcodes are read out by sequential hybridization, similar to ORCA barcodes (**Fig. 1b, 1c**). This approach both accelerates data collection and removes potential batch effects, intrinsic to high-content, high-resolution assays. After imaging, we demultiplex cell labels and assign the single chromosome traces from ORCA to individual cells from each genotype and each treatment condition. In the multiplexed cell image shown in Fig. 1C, we combined the parental E14 cell line with RAD21 and CTCF lines +/- auxin treatment. Replicates showed high reproducibility (**Extended Data Fig. 1b**). To increase cell numbers and statistical power, these were merged for all subsequent plots and analyses.

We can compute population-level contact frequency maps with ORCA by calculating, for each pairwise combination, the fraction of chromosomes that are within a certain distance (we used 200 nm, approximately equal to the mean distance between the centers of adjacent regions). Plotting contact frequency over all ORCA traces in each condition (**Fig. 1d**) we found good agreement with Hi-C contact maps (**Extended Data Fig. 2a, b**). Cohesin-depleted contact maps showed loss of TADs ^17,18^ in CTCF-depleted cells, TADs were still visible, although with lower insulation between them (**Fig. 1d**, **Extended Data Fig. 2c**). Published Hi-C data of CTCF depletion in mESCs ^16,19^ also showed that, unlike cohesin depletion ^18^ CTCF loss did not fully abolish TADs for these regions (**Extended Data Fig. 2a, b**).

Thus, with the ability to 1) rapidly deplete CTCF and cohesin, 2) multiplex multiple cell types and treatment conditions, and 3) measure genome structure that matches well at the population level with Hi-C, we set out to examine in detail how these proteins affect genome folding in the subsequent sections.

### CTCF contributes to TAD separation while enabling cross-TAD contacts by organizing chromosomes radially

In ORCA and Hi-C population contact frequency maps for unperturbed cells, we observed loops that span CTCF-marked TAD borders, which we called ‘bridges’ for short (**Fig. 2a, b, Extended Data Fig. 3a, b**). Hi-C data show such bridges throughout the genome (e.g. ^29^), though the population-average nature of the data does not distinguish between multiple structural origins of this feature. Such a bridging pattern in contact frequency could arise from a heterogeneous population in which distinct portions of the population have either the long-range contact or the separated structure (**Fig. 2a**). To test for this, we turned to our single-chromosome data and examined the all-to-all contact frequency computed using only those chromosomes which contained bridging contacts (**Fig. 2b, Extended Data Fig. 3c**). Interestingly, this revealed that the population of chromosomes with bridging contacts still contained clear TAD borders beneath the bridges (**Fig. 2b, Fig. Extended Data Fig. 3c**). Thus, a bridge can occur between separated structures and the two scenarios (**Fig. 2a**) are not mutually exclusive.

**Figure 2.**
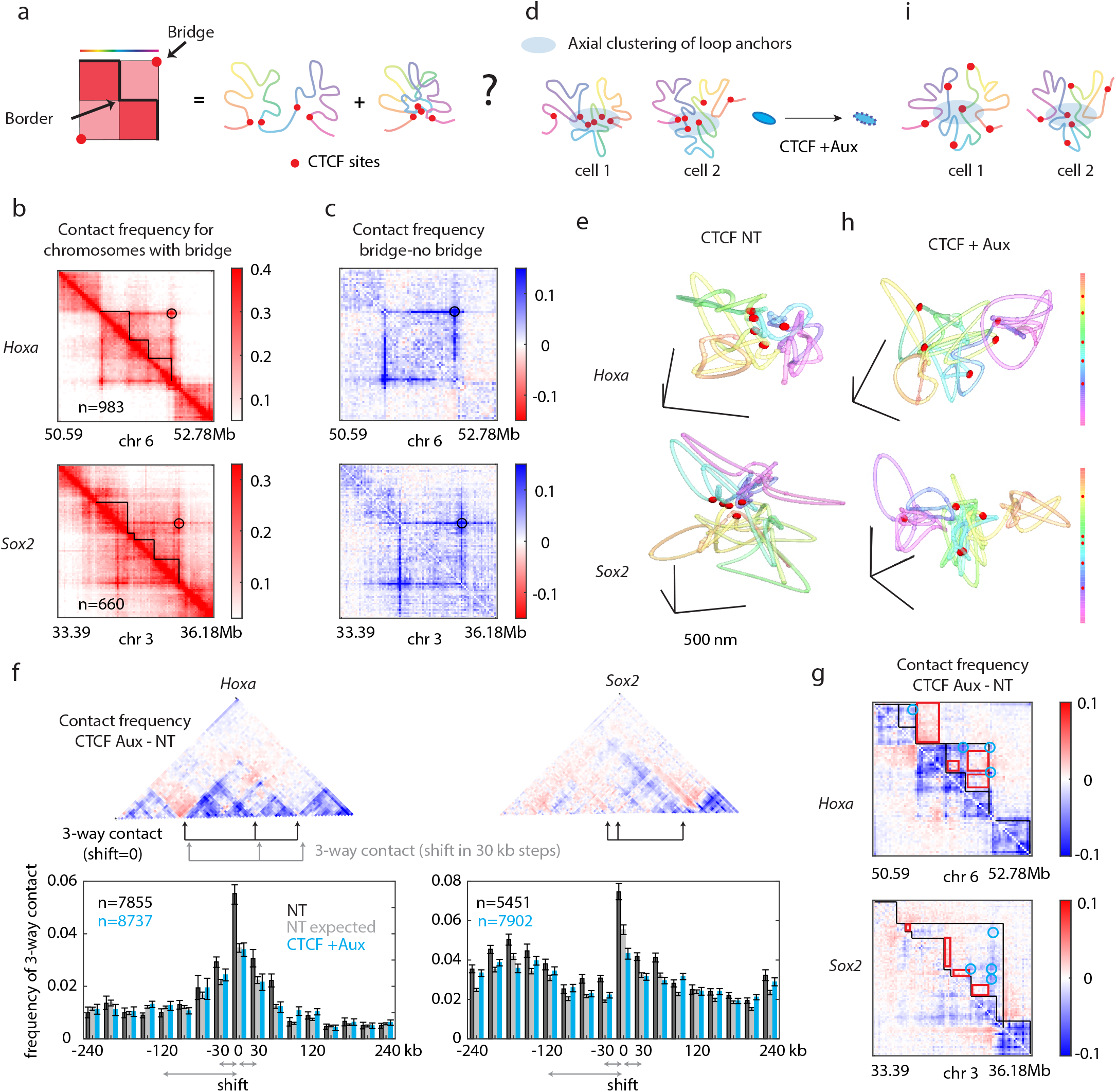
CTCF enables cross-TAD interactions. **a.** Schematic depiction of bridges (left) and schematic hypothesis how these arise from a mixed population containing either the bridge contact or the domain separation. **b**. Contact frequency map from all cells with the bridging contact indicated (black circle). **c**. Subtraction of contact frequency maps from all cells that have the indicated bridging contact minus those without it. **d**. Schematic depiction showing how CTCF interactions create a radially organized chromosome. Near the center axis (gray region) CTCF sites (red) and CTCF-proximal regions make multi-way interactions. **e**. Examples of single chromosomes with CTCF-marked boundary positions colored in red. **f**. Top panels, contact frequency difference map between CTCF Aux and untreated cells, with arrows indicating the positions chosen for the 3-way analysis. Lower panels: Frequency of 3-way contact between the three CTCF-marked positions indicated by black arrows. Frequency of 3-way contact for other distance-matched triplets (black) is shown as a function of how far (in kb) they are shifted from the black triplet. In gray, expected 3-way contact based on the pairwise contact probabilities and in blue, frequencies of 3-way contact for CTCF auxin-treated condition. Error bars show confidence intervals (see Methods). **g**. Contact frequency difference map for CTCF auxin-treated and untreated cells. Black lines show TAD boundaries. **h**. Example chromosome traces for CTCF auxin-treated cells. **I**. Schematic illustrating the effect of CTCF loss on randomizing the position of CTCF-marked boundary sites.

To better see how these bridge chromosomes differ from non-bridge chromosomes we subtracted their contact maps. The difference maps revealed a prominent stripe connecting the bridge anchors, with strengthened loops among intervening CTCF sites in bridge-containing chromosomes (**Fig. 2c**, **Extended Data Fig. 3d**). This implies that bridge anchors frequently contact several additional sites in between, with a preference for intervening CTCF sites, rather than spanning anchor-to-anchor in a single loop (**Fig. 2d**). Inspection of individual chromosome traces revealed examples of distal CTCF sites bridged by multiple loops, including intervening CTCF sites (**Fig. 2e**).

We next computed the frequency of three-way contacts among CTCF sites. We compared these to distance-matched, non-CTCF interactions using a sliding window across the genomic region. Three-way contacts were more common among the CTCF sites than non-CTCF sites (**Fig. 2f**). Notably, CTCF-site-proximal regions also showed elevated rates of three-way contacts, which fell off as a function of distance from the CTCF-site (**Fig. 2f**). The three-way contact frequency among CTCF sites was also higher than expected based on the measured pairwise interaction frequencies (assuming pairs are independent), providing further evidence of preferential clustering missed in bulk pairwise data (**Fig. 2f**). Depletion of CTCF significantly reduced the frequency of three-way interactions among both CTCF and CTCF proximal sites, though made little change for CTCF-distal sites (**Fig. 2f**).

To explore the effects of CTCF loss across the domain, we computed the change in contact frequency upon CTCF degradation. We note that ORCA contact frequency is in absolute terms (fraction of chromosomes), rather than normalized within the experiment, enabling a direct subtraction between contact frequencies. Previous microscopy studies ^23,24^ and simulations ^21,30^ concluded that the distance between TADs decreased upon CTCF depletion. ORCA data from the *Hoxa* and *Sox2* domains revealed a more complex pattern upon CTCF depletion, where some pairwise distances increased while others decreased (**Fig. 2g**). In particular, while several of the CTCF borders exhibited increased interaction between the upstream and downstream TADs, TAD bridges exhibited reduced interaction, even though the contact frequency increased between TADs underneath the bridge (**Fig. 2g** red box, blue circle).

Together these single-cell analyses point to a tendency of CTCF to organize chromatin in a radial pattern, rather than functioning purely as a border between domains or a tether within a domain (**Fig. 2d**). CTCF-border sites preferentially position on the axial part of this structure (**Fig. 2d** gray region), where they are more likely to interact, including across boundaries. Intra-TAD regions (far from CTCF sites) preferentially position away from this axis, where they are generally more isolated from other intra-TADs. Depletion of CTCF reduced this clustering tendency of the CTCF-marked borders and led to more disordered positioning of loops (**Fig. 2h, i**), which led to a weakening of TAD borders on the population-level contact frequency maps (**Fig. 1d**). We note that although this radial organization is reminiscent of the rosette, single-cell traces show considerable heterogeneity of the number and identity of sequences at the center, and a relatively low frequency of multi-way contacts for any particular triplet, even though triplet interactions among CTCF sites are substantially more common than non-CTCF sites (**Fig. 2f**).

### Cohesin loss increases distances both within and between TADs

We next examined the role of cohesin in this radial chromosome organization. Degradation of cohesin clearly reduced the preference for CTCF and CTCF-proximal regions to form three-way contacts (**Extended Data Fig. 3e**). In contrast to CTCF depletion, three-way contacts among all loci decreased, both CTCF-proximal and CTCF distal (**Extended Data Fig. 3e**). Inspecting individual chromosome traces, we found that depletion of cohesin led to less loopy structures and expansion of the traces (**Fig. 3a**), unlike CTCF, where loops persisted in CTCF-depleted cells but with more random positioning of TAD borders (**Fig. 2h, i**).

**Figure 3.**
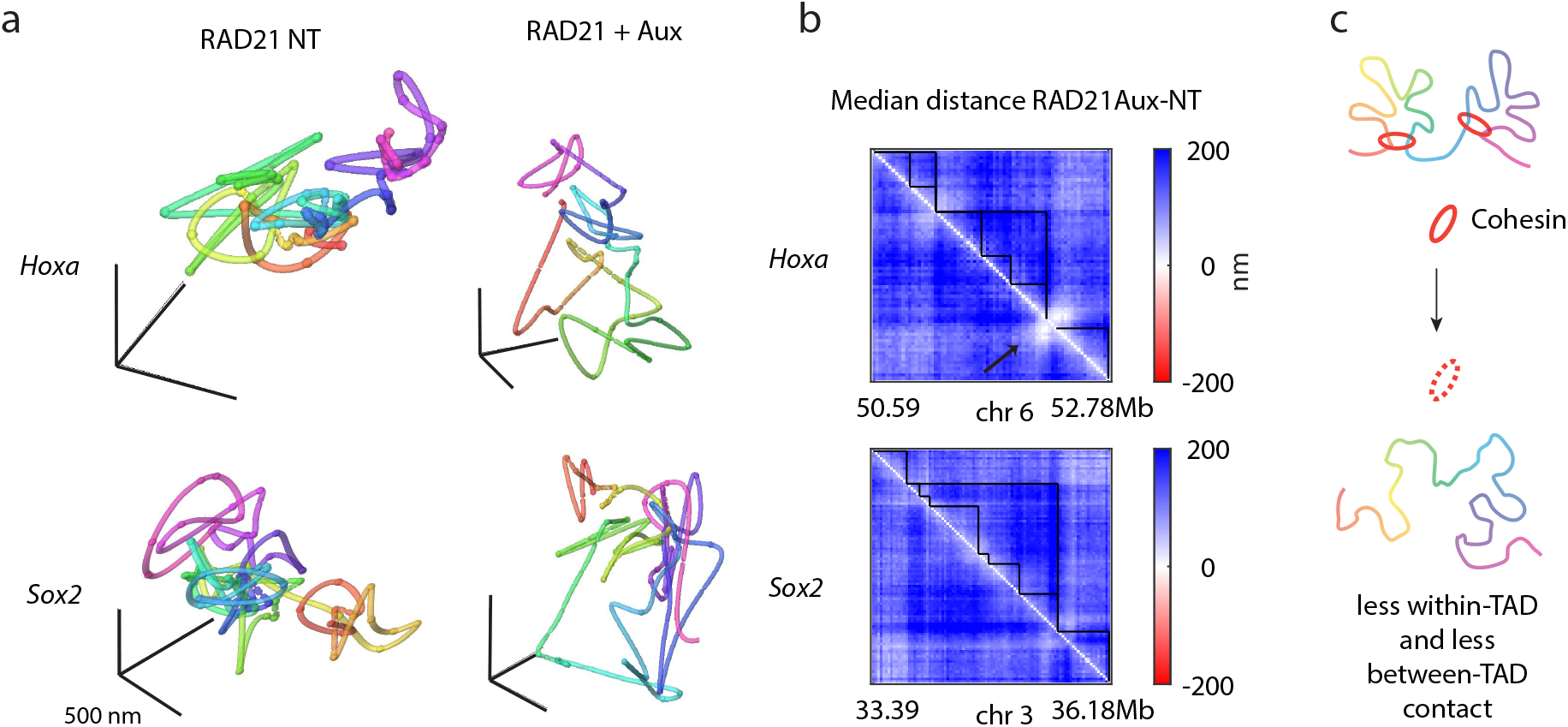
Cohesin loss leads to increased distances within and between TADs. **a.** Example single chromosome traces for *Hoxa* and *Sox2* regions from untreated and auxin-treated cells. **b**. Differences in median distance between auxin-treated and untreated cells show that all pairwise distances increase (all blue). TAD outlines are drawn in black. For the *Hoxa* region, the area indicated by the arrow corresponds to the *Hoxa* gene region that is coated by Polycomb and does not expand upon auxin treatment (white). **c**. Schematic illustrating that loss of cohesin leads to general expansion, increasing distances within and between TADs.

To quantify the expansion after cohesin depletion across the whole population, we plotted the median pairwise distances in units of nanometers (**Extended Data Fig. 4a**). We found that depletion of cohesin led to increased pairwise distances by up to a few hundred nanometers for both *Hoxa* (2.2 Mb) and *Sox2* (2.8 Mb) domains (**Fig. 3b, Extended Data Fig. 4a**). Distances increased significantly (ranksum, *p*<0.05) **(Extended Data Fig. 4b**) for all pairwise interactions both within and between TADs, with the fold change of expansion being greater within TADs (**Extended Data Fig. 4c**). The only non-significantly changed interactions (**Extended Data Fig. 4b**) we observed were at the Polycomb-bound *Hoxa* region (**Fig. 3b, Extended Data Fig. 4d arrow**) which remained compact and maintained contact with another upstream Polycomb-bound region in 19.4% of cells (relative to 29.7% in untreated cells) (**Extended Data Fig. 4d**).

An advantage of the multiplexed microscopy approach is that changes in distance are measured in absolute units of nanometers, and systematic batch effects between experiments are removed by analyzing both treated and untreated cells at the same time. It is more challenging to make direct comparisons between treatment conditions with Hi-C data, as there are no spike-in control methods available to enable normalization across experiments. This uncertainty may explain why the untreated/cohesin-depleted ratio of Hi-C maps for *Hoxa* and *Sox2* suggested reduced contact within and increased contact between TADs (**Extended Data Fig. 4e**), in contrast to the ORCA data - even though the original, non-subtracted contact frequency maps agree qualitatively for both conditions (**Fig. 1d**, **Extended Data Fig. 2b**).

To test whether this expansion was dependent in part or in full on the loss of sister chromatid cohesion, we performed immunofluorescence labeling of Geminin and ORCA in the same cells (**Extended Data Fig. 4f**). Geminin is a cell-cycle-regulated protein that is not detected in G1 and progressively accumulates through the S and G2 phases of the cell cycle. We selected cells with low Geminin levels (lowest 25% of the cells) and found that these also showed expansion upon loss of cohesin, both within and between TADs (**Extended Data Fig. 4g**). Further, we found no correlation between Geminin levels and the median pairwise separation among all imaged domains on a single-cell basis (**Extended Data Fig. 4h**). From this we conclude that the expansion of chromosomes at the megabase scale is primarily driven by changes in cis-structure, and not loss of sister cohesion.

In summary, not only do we find that TAD loss is the consequence of increased separation of sequences within the TAD (and not mixing between TADs), but that it occurs in the context of a significant general expansion of the genome at the megabase scale (TAD scale) (**Fig. 3c**). Our findings are consistent with previous measurements at this scale that found an increased separation of genomic loci in *cis* upon cohesin loss in fixed cells ^20,23,24,31–34^ and live cells ^35,36^.

### Cohesin loss leads to increased mixing at the chromosome scale

Increase of pairwise distances along the ∼2.5 Mb scale upon loss of cohesin made us wonder what the consequences were on genome organization at the nuclear scale. We first quantified the nuclear area among untreated and cohesin-depleted cells as well as the E14 parental cell line. We observed a similar distribution of nuclear sizes in all conditions (**Extended Data Fig. 5a**).

Given the expansion of distances at the 2-3 Mb scale and lack of change in nuclear size, we wanted to understand what happens to the genome at the tens-of-megabases and whole-chromosome scales. We designed probes spanning both a ∼30 Mb region and the whole chromosome (∼150 Mb). We did this for both chromosomes 3 and 6 that contain the *Sox2* and *Hoxa* domains examined previously (**Fig. 4a**).

**Figure 4.**
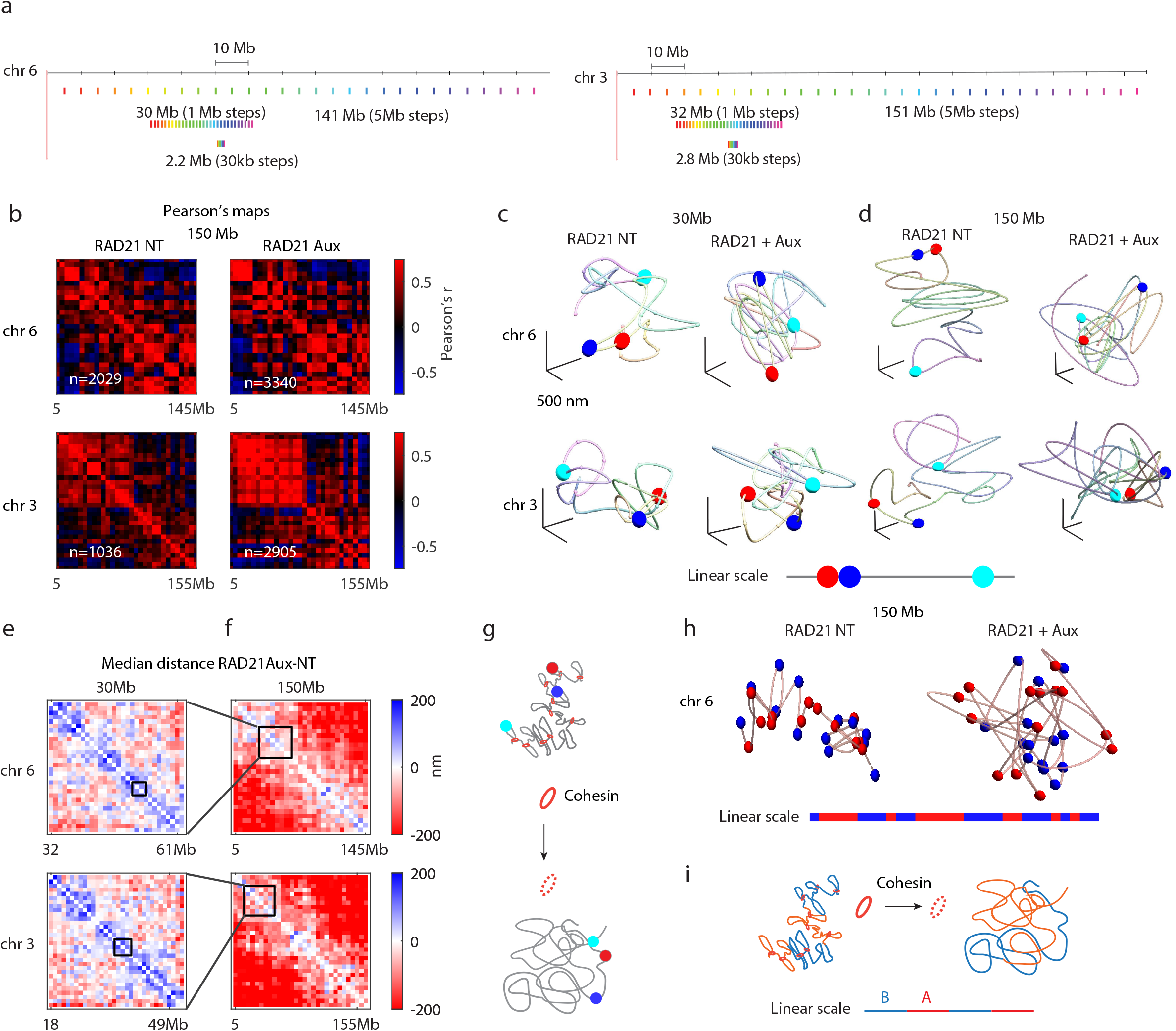
Cohesin loss leads to increased mixing at the chromosome scale. **a.** Schematic of probes designed to image folding of ∼30 Mb and ∼150 Mb regions of the genome, shown relative to the ∼2.5 Mb probes imaged previously. **b.** ORCA Pearson correlation maps for the 150-Mb scale probes in RAD21 untreated or auxin-treated cells. **c-d.** Examples of single chromosome traces for ∼30 Mb (c) or ∼150 Mb (d) regions of chromosome 6 and chromosome 3 with or without auxin. Probes that correspond to consecutive (red and blue) and a more distal (cyan) genomic distances are represented with larger spheres. Their interprobe distances are shown in the linear colorbar. **e-f**. Differences in median distance between RAD21 auxin-treated and untreated cells for the ∼30 Mb (e) and ∼150 Mb (f) regions, show that distal genomic regions are closer (red) after auxin treatment. The black squares in (e) show the position of the ∼2.5 Mb domains imaged previously, while the black squares in (f) show the position of the ∼30 Mb domains. **g.** Schematic illustrating the disordered chromosome structure upon cohesin degradation. **h.** Example chromosome traces in untreated or RAD21 depleted cells colored by compartment identity, as in (b). **i**. Schematic illustrating the effect of cohesin degradation on compartments.

Previous work using Hi-C found that depletion of cohesin did not significantly alter the chromosome-scale genome organization other than a slight strengthening of compartments ^17,37^. Compartments refer to a plaid pattern commonly seen at the chromosome scale in Hi-C contact maps, indicating a preferential separation of the genome into two groups, commonly denoted A and B ^38^. The separation is readily visualized through the Pearson map, where each element of the map reports the cross-correlation of the corresponding row and column after distance normalization ^38^. Examining the Pearson maps for the 30-Mb and full-chromosome ORCA data, we observed a similar plaid pattern to Hi-C (**Fig. 4b**, **Extended Data Fig. 5b, c**), an agreement also observed in compartment analysis from earlier tracing experiments ^39^. Upon cohesin depletion, we also observed a slightly stronger plaid pattern and higher Pearson’s r, indicating compartment strengthening (**Fig. 4b, Extended Data Fig. 5b-e**).

We then looked at single-chromosome traces to understand how the expansion we observed at the 2-3 Mb scale could be reconciled with minor compartment changes at the 30 Mb and 150 Mb scales. The 30-Mb and whole-chromosome traces looked more disordered after depletion of cohesin. While in untreated cells, adjacent labels (**Fig. 4c, d** red and blue labels) remained in physical proximity relative to genomically distant regions (red and cyan), in cohesin-depleted cells, there was less distinction in terms of physical distance between genomically proximal or distal labels. To test these effects population wide, we plotted the difference of median distance between auxin-treated and untreated cells for all pairwise combinations (**Fig. 4e, f**). We found that at the 30-Mb scale (**Fig. 4e**), pairwise distances less than ∼4 Mb apart, increased significantly (**Fig. 4e, Extended Data Fig. 5f**), consistent with our data showing expansion at the TAD scale (**Fig. 3a, b**). However beyond the ∼4 Mb separation we found a gradual transition from increased to decreased distance (**Fig. 4e**, blue to white to red). The decrease in distance upon cohesin depletion became more and more pronounced and significant (**Extended Data Fig. 5f**) as the genomic distance between two pairs increased, as seen in the chromosome-scale difference maps (**Fig. 4f**).

Our ORCA measurements at the 30-Mb and chromosomal scale show that cohesin is essential for maintaining an ordered chromosome structure (**Fig. 4g**). By primarily contributing to a more linear ordering of the chromosome, cohesin also opposes the demixing of compartment types (**Fig. 4h, i**), explaining the slight compartment strengthening after cohesin depletion.

### Loop extrusion simulations can exhibit multiscale effects on genome folding

Considerable evidence supports the model that cohesin shapes genome structure, including TADs, through loop extrusion (for review see ^2,40^). In this model, cohesin is loaded onto DNA and extrudes a loop until it is stalled by CTCF. We wanted to know if our ORCA observations of expansion at the ∼2.5 Mb scale (**Fig. 3**) and increased mixing at the 30-Mb and full-chromosome scales (**Fig. 4**) could also be understood through the process of loop extrusion. We performed Langevin dynamic simulations of loop extrusion extended from previous polymer models ^21,30^ (**Methods**). As expected ^21^, simulations including cohesin formed TADs between CTCF boundaries, and loops between convergent CTCF sites, while simulations without cohesin did not **(Extended Data Fig. 6a**). Examining individual traces at the 3-Mb scale, we found that without cohesin loops, the polymer trace expanded (**Fig. 5a**). To follow structural changes at the chromosome scale, we plotted the whole chromosome structure and colored 3-Mb-sized regions blue, red and cyan (**Fig. 5b**). With cohesin, adjacent regions (blue and red) could be found proximal in 3D space (**Fig. 5b**). In contrast, without cohesin, individual regions expanded significantly, such that the ends of a 3-Mb region could be near opposite sides of the chromosome territory (e.g. red domain and gray chromosome, **Fig. 5b**). This expansion also facilitated proximity between linearly distal elements in some chromosomes (**Fig. 5b**, blue and cyan regions). This expansion at shorter scales and contraction at larger scales could be seen across the population of replicate simulations, as shown by the difference of the median distance maps (**Fig. 5c, and Extended Data Fig. 6b**. Thus, cohesin-mediated loops reduce disorder in chromatin folding by ensuring linearly proximal regions remain proximal in 3D and linearly distal regions remain distal in 3D.

**Fig. 5.**
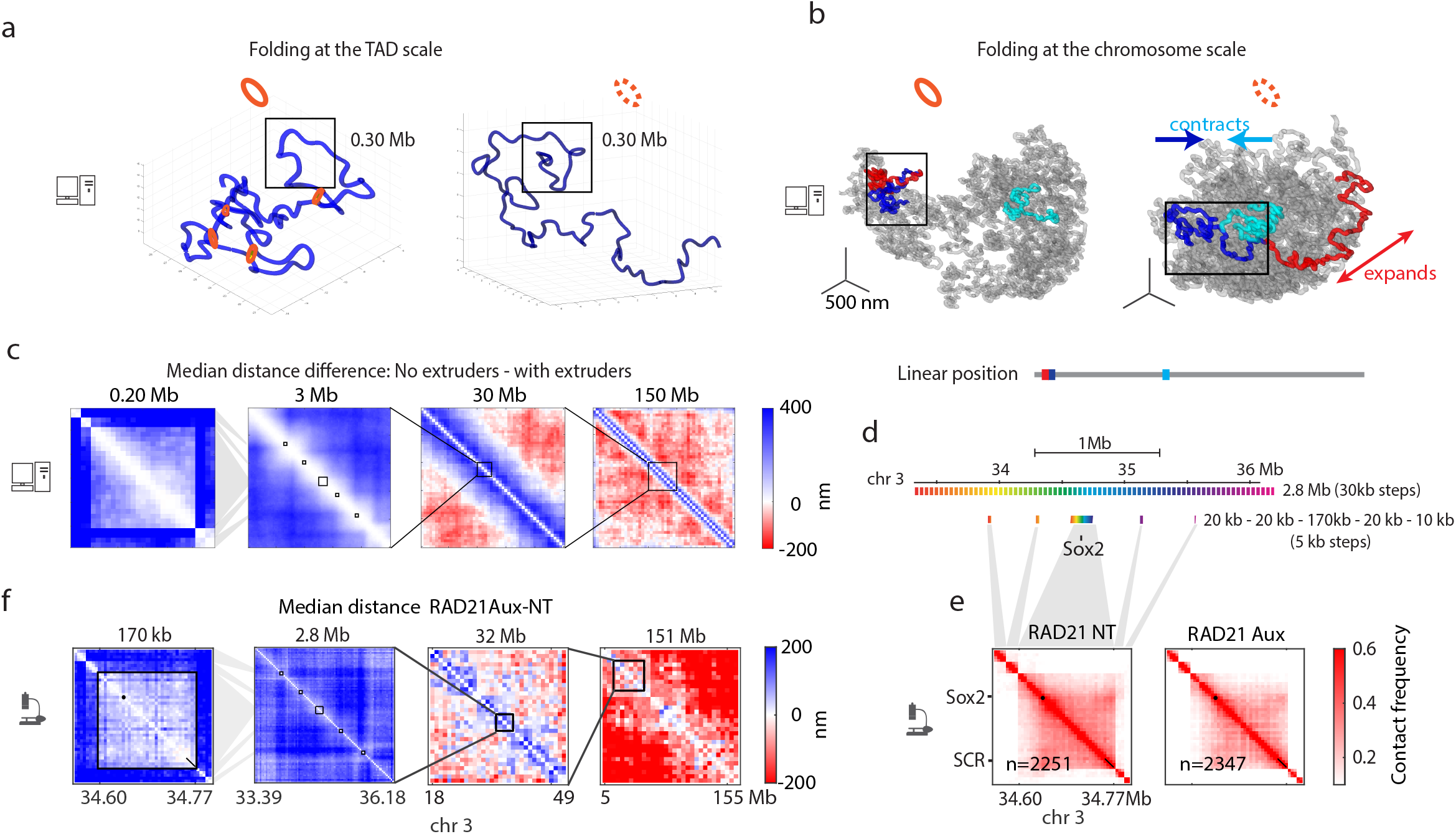
Loop extrusion model shows the multi-scale effects of cohesin loss. **a.** Simulated polymer corresponding to 3 Mb in length with or without loop extrusion factors. The regions shown are a zoom-in view of the blue region inside the black boxes in (b). **b**. Polymer simulations at the chromosome scale (150 Mb) with or without loop extrusion factors. The chromosome is shown in gray. Other chromosomes inside the simulated nucleus are not shown to avoid overlap. Individual 3 Mb domains are colored. Their positions on the linear polymer are indicated by the colorbar. **c.** Difference in median distance between simulations with and without loop extrusion at different genomic scales: 3 Mb, 30 Mb and 150Mb. Zoom-out of the 3 Mb corresponds to a 200 kb domain and a few more distally spaced regions. **d**. Schematic of the probe design for a 170 kb region encompassing the *Sox2* gene with probes spaced 5 kb apart, as well as 4 adjacent and more distally spaced regions, relative to the 2.8 Mb probeset used in Figs. 1-3. **e.** ORCA contact frequency maps for finer-scale Sox2 probes in RAD21 untreated or auxin-treated cells. **f.** Difference in median distances between RAD21 auxin-treated and untreated cells for the 2.8-Mb sized domain (same as in Fig. 3b and Fig. 4e, f) and the finer-scale region. Squares on the 2.8 Mb map indicate the regions that were imaged at higher resolution.

Not all parameter choices for loop extrusion produced the switch from expansion to contraction that we observed in our experimental data and in the simulations described above. Simulations with low-density confinement (confined to 10% of their relaxed volume) exhibited increased separation at all length scales (**Extended Data Fig. 6c**). Chromosomes with a higher frequency of chain-crossing behavior (as might be expected for a high level of topoisomerase, see **Methods**) also failed to exhibit the qualitative switch, expanding at short scales and exhibiting no change at larger scales (**Extended Data Fig. 6d**). Thus, only with substantial confinement and reduced chain-crossing did the cohesin-loops significantly hold apart the more distal parts of the chromosome by steric exclusion – loop-clash. This finding is consistent with recent polymer theory showing that chromatin loops “experience topological repulsion” ^41^.

Furthermore, our simulations predict that at length scales less than ∼200 kb, the relative effect of cohesin loss on 3D contacts would be less than observed at the Mb scale (**Fig. 5c**), potentially because 200 kb is shorter than the typical distance between cohesins (**Fig. 5a**, black boxed regions). To test if <200 kb regions are indeed less affected by cohesin loss, we labeled at a higher resolution (5-kb step size) the 170 kb region containing the Sox2 gene and its downstream enhancer ^42,43^, as well as more distal regions flanking this domain, 200 kb to 1.5 Mb apart (**Fig. 5d, e**). Upon depletion of cohesin, we found that within the 170 kb region, expansion was significantly lower (ranksum p=1.03*10-168) than for the more distally spaced probes (**Fig. 5f, Extended Data Fig. 7**).

Thus we find that the loop extrusion model is consistent with ORCA data collected at all scales (**Fig. 5c, f**), if sufficiently confined in the nucleus and sufficiently restricted in the ability for chain-crossing. The model does not quantitatively match the data in all respects, but does show how loop extrusion, with a few parameter constraints, can explain the key qualitative trends in the data. The picture that emerges is that extrusion of chromatin loops by cohesin locally compacts DNA both within TADs and across TAD borders, and that these loops further serve to organize the chromosome by sterically inhibiting longer-range contacts that would otherwise be promoted by the dense packaging environment of the nucleus.

### Cohesin depletion increases the variability in gene expression

Previous studies have shown that degradation of cohesin has a limited immediate effect on transcription ^17,18,44–46^ relative to degradation of transcription factors or co-factors ^47–50^ or genetic disruptions of TAD borders ^5,51–54^. Given the increased disorder in genome structure, where proximal regions got farther apart and distal genomic regions became closer, we wondered if at the single-cell level cells would exhibit increased variability in gene expression.

We performed multiplexed single-molecule FISH to measure expression of a set of developmentally regulated pluripotency genes. We used cellular barcoding to minimize batch effects among different treatment conditions and cell lines (RAD21-AID, CTCF-AID, E14) (**Fig. 6a**). We also added mock (DMSO) treated cells as a further comparison for the magnitude of the effect. The mean mRNA counts showed small fold changes between treatment conditions. Effects of auxin on median depletion were scarcely stronger than for our mock treatment (**Fig. 6b**). To see if cohesin depletion led to increased variability between cells, we calculated the coefficient of variation. We found that the majority of genes tested showed a modestly higher coefficient of variation after cohesin depletion (**Fig. 6c**). Depletion of CTCF had less effect on the coefficient of variation (**Fig. 6c**), likely reflecting less change in genome structure after CTCF depletion at either the TAD scale or chromosome scale (**Extended Data Fig. 8**).

**Figure 6.**
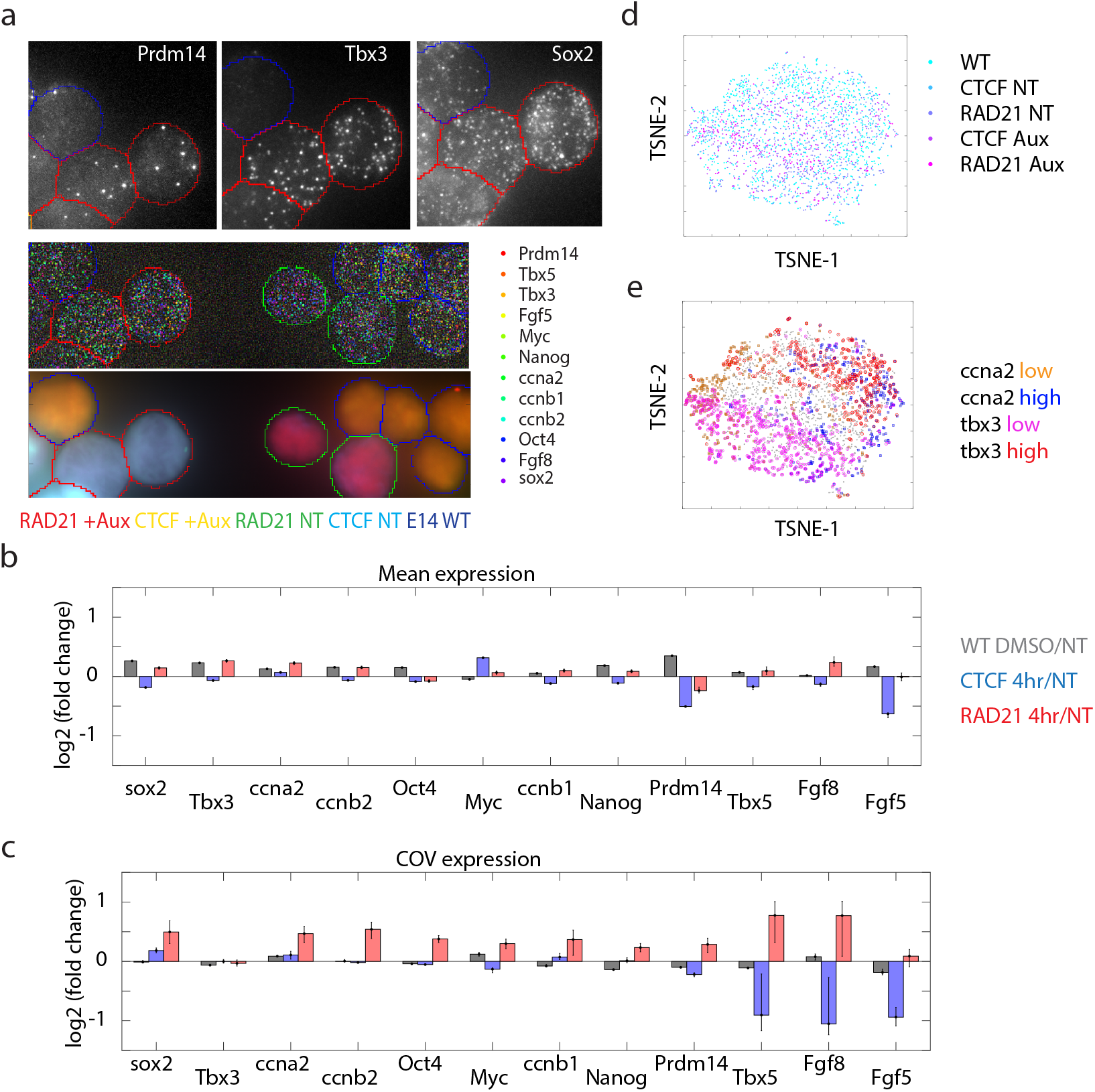
Cohesin depletion increases the variability in gene expression. **a.** Example of multiplexed cells for different treatment conditions labeled by single molecule FISH. **b**. Mean fold change of expression. **c**. Fold change in coefficient of variation (COV).

Cluster analysis by TSNE showed that RAD21-depleted cells did not split into distinct groups, nor cluster separately from untreated cells (**Fig. 6d**). This suggests increased variability was not due to partial differentiation of the mESCs after auxin treatment for 4 hours. Indeed all the genetic backgrounds and treatment conditions clustered together (**Fig. 6d**), with variation of cell-cycle-dependent factors aligning with one major TSNE axis and variation in *Tbx3* defining the other (**Fig. 6e**), a gene which showed little CTCF or cohesin dependent change (**Fig. 6b**).

Overall our data show that acute loss of cohesin leads to disordered chromosome structures that result in modestly increased cell-to-cell variability in gene expression, which is missed by bulk population assays.

## Discussion

Here we introduced a multiplexed approach for ORCA, enabling simultaneous measurement of chromosome folding across multiple treatment conditions. Our approach enables quantification of physical structure in absolute units, enabling quantitative comparison between conditions, overcoming a limitation of Hi-C. As the field moves increasingly to the study of perturbations, we expect the ability to measure absolute frequency differences across genomic length scales without normalization artifacts or batch effects, the ability to measure variation in single cells, and the ability to process multiple conditions at a time will facilitate more rapid understanding. In this work, we characterized the structure of the interphase genome from the kilobase to the chromosome scale, expanding our understanding of the role of CTCF and cohesin in shaping that structure, with implications for gene regulation.

CTCF is thought to determine TAD borders by defining the bounds of cohesin-mediated loop extrusion ^1–4^. Consistent with this, 2-color FISH experiments ^19,23,24^ reported a decrease in distances between TADs, and an increase in overlap. While supporting these conclusions, our ORCA data revealed more complex patterns upon CTCF loss. In addition to supporting TAD boundaries, we identified a role for CTCF in bridging across TAD borders through a multi-loop structure. This behavior provides a mechanism that would enable contacts between enhancers and promoters that are separated by one or more TADs. Indeed, one of the CTCF-border-crossing sites in the *Hoxa* locus has recently been shown to contain enhancer activity for *Hoxa* genes during hematopoiesis ^55^ and cranial-facial patterning ^56^. Furthermore, several groups recently inserted CTCF sites between the *Sox2* gene and its embryonic enhancer region, but found minimal impact on *Sox2* expression, despite establishing a new TAD border ^57,58^. As both the enhancer region and promoter are each already CTCF proximal, the multi-way clustering of CTCF could explain this insulator bypass as well.

The radial organization arising from the preferential multi-way contacts of CTCF sites resembles earlier models of chromatin structure based on 3C assays, like the archipelago of enhancers and promoters described for *Hoxd* ^59^, the subTAD-clusters described for *Hoxa* ^60^, or the CTCF hubs described for the *protocadherin* genes ^61^, or B-globin genes ^62,63^ among other regions of the genome ^61,64^ Multi-contact 4C and multi-contact-Hi-C experiments have provided further evidence for three-way interactions among CTCF sites ^63,65^. By observing the structure of individual chromosomes, we provide more direct evidence for these multi-way interactions and how they maintain intervening borders. We also see they exist within a heterogeneous population, in which diverse parts of the chromatin fiber alternate at loop bases and that any particular triplet is individually rare.

Furthermore, CTCF degradation, compared to cohesin degradation, exhibited substantially weaker impacts on TADs in our data, consistent with recent reports ^16,19,66^. Whether this is due to co-factors at some sites which help retain CTCF locally amidst global depletion or additional factors contributing to cohesin-dependent boundary activity ^57,67^ remains an interesting question for future studies.

Cohesin depletion led to a disordered genome structure with distinct effects at different genomic scales. At the ∼200 kb scale, typical of the majority of enhancer-promoter interactions ^68^, we found relatively mild increases in pairwise distances as compared to the ∼2.5 Mb scale which showed a significant expansion after depletion of cohesin. Thus, effects of cohesin loss would be different for each gene. Genes whose enhancers are within ∼200 kb distance, will be less affected in terms of E-P contact than genes that rely on more distal enhancers (>200 kb - 3 Mb away). At larger genomic scales (>4 Mb) we found increased mixing. This implies that all genes are more likely to experience contacts with very distal enhancers (>4 Mb to 150 Mb away) and these ectopic enhancer-promoter contacts will be different for each cell. While any particular contact occurs at relatively low frequency, each promoter is now exposed to many more such rare long-range contacts than before. With this perspective, rather than expecting a global up- or down-regulation of genes after depletion of cohesin, our ORCA measurements suggested that each gene and each cell would behave differently. This would manifest as increased cell-cell variation in mRNA levels, which we observed for most of the developmentally important genes we assayed by single molecule RNA FISH. Recent studies have similarly reported that disruptions in chromatin 3D organization increase variability in gene expression, using single-cell RNA-seq ^69^, RNA-FISH and flow-cytometry ^70^, or integrative modeling of omics data ^71^. Whether and how these acute changes in cell-cell variability due to disruption of chromatin structure connect to functional outcomes of cohesin loss or reduction in development and disease ^72–77^ will require future investigation.

The structural changes we observe in genome organization at the chromosome scale also allow us to revisit recent conclusions about the role of cohesin in aspects of genome organization, such as compartments and polycomb loops. It has been proposed that cohesin opposes demixing of the genome into A/B compartments directly, through its ability to bridge compartment boundaries and pull together A and B chromatin ^17,30,37^. This proposal is supported by the observed strengthening of A/B compartments upon cohesin depletion ^17,37^. Our data suggest an additional mechanism. By significantly decreasing the frequency of interactions >4 Mb, through the dense stacking and clash of loops, cohesin indirectly disrupts compartmentalization by reducing the opportunity of linearly separated A (or B) domains to contact and mix. While both the direct and indirect effects of cohesin may operate in tandem, the direct explanation predicts that short-range compartmentalization should be most affected (where A meets B along the linear genome), whereas the indirect mechanism suggests long-range interactions should be most affected.

It has been proposed that cohesin processivity can disrupt the Polycomb (Pc) homotypic interactions which drive formation of ultra-long range contacts among Polycomb domains (1-20+ Mb), “Polycomb loops” ^18^. This data is supported by the observation that these loops strengthen upon cohesin degradation ^18^. As we have shown for the 1-Mb scale interaction in the *Hoxa* region, the shorter Pc-Pc interactions actually decrease in absolute frequency upon cohesin loss, while >4 Mb interactions increase along the entire chromosome. Thus, the effect of cohesin on Pc loops we propose is also an indirect effect of cohesin linearizing the chromosome (promoting short <4 Mb contacts and reducing longer range ones). Indeed, shorter range Pc loops were also reported to show less change or increase interaction by Hi-C ^18^, an effect readily explained by the loop-clash model that would not be expected if cohesin linear scanning were to knock apart Pc-Pc loops.

Together, these data suggest cohesin and CTCF organize the genome through their distinct effects on chromatin loops and loop stacking. Cohesin increases the density of chromatin loops, and thus proximal sequences are brought into frequent contact by individual loops and stacking loops, while distal sequences are held apart by the clash of many loops. When loops stack together, the loop bases aggregate in a central hub. By stalling cohesin, CTCF sites are more likely to be found at loop bases, and more likely to end up at the hub. This aggregation of 3 or more borders facilitates cross-border contacts among CTCF-proximal regions, while maintaining border function among CTCF-distal regions. Thus, beyond its known functions in enhancer blocking and looping, CTCF may also facilitate cross-TAD gene regulation.

## Methods

### Cell culture, treatment and collection

E14 parental cell line as well as the Rad21-AID and CTCF-AID cell lines were a gift from Elphege Nora. Cells were grown in DMEM/F12 supplemented with 15% FBS, NEAA, Glutamax. A final concentration of 500uM IAA (Sigma i2886) was used for all auxin treatments. Cells were treated for 4 hours, media was changed in the untreated condition at the same time as the fresh media with auxin was added to auxin treated cells.

For cell collection, cells were trypsinized, counted, spun down and after removal of supernatant media, were fixed in suspension in 4%PFA in 1xPBS on a nutator for 10 minutes, washed 3 times with 1xPBS, resuspended in 70%EtOH in 1xPBS at a concentration of 1-10 000 cells/ul and stored at -20C for up to 6 months.

### Western blot

Cells were collected by trypsinization, washed in 1xPBS and pellets were frozen on dry ice/EtOH bath and stored at -80C. Cells were lysed in mRIPA buffer with Protease and Phosphatase inhibitor cocktail. 20ug of protein was run on a 4-20% Mini-PORTEAN TGX precast gels in Tris-Glycine-SDS running buffer. After transfer onto the nitrocellulose membrane, and blocking in Li-Cor Intercept Blocking Buffer (TBS), blots were probed with either CTCF (Abcam 128873, 1:1000) or Rad21 (Abcam 154769, 1:1000) antibodies as well as beta-Actin (Cell Signaling 3700) in blocking buffer at 4C overnight. Membranes were then washed 3x 5min in TBST at room temperature, incubated with secondary antibodies (Li-Cor: IRDye 680RD Donkey anti-Rabbit 926-68073 and IRDye 800CW Donkey anti-Mouse, 1:10 000 diluted in blocking buffer) for 1h at room temperature. After 3x 5min washes in TBST, and 1 wash in TBS, samples were imaged on the Li-Cor scanner.

### Multiplexed cell labeling

The multiplexed cell labeling protocol was adapted from Gehring et al. 2020. The detailed protocol is as follows.

#### Part 1: Preparation of labelled oligos

(1) 3’ amine-modifed oligos were ordered from IDT and resuspended to 500 µM in 50 mM sodium borate buffer pH 8.5
(2) In 1.5 mL microcentrifuge tube, add 25.0 µL 3’ amine-modified oligo in 50 mM sodium borate buffer, 8.2 µL 10 mM Methyltetrazine-NHS ester, 41.8 µL DMSO, incubate protected from light for 30 minutes with agitation
(3) After 30 minutes, add another 8.2 µL 10 mM Methyltetrazine-NHS ester, then add once again after an additional 30 minutes
(4) After the 90-minute reaction, quench by adding 180 µL 50 mM sodium borate buffer and 30 µL 3 M NaCl, then add 750 µL ice cold 100% EtOH]
(5) Precipitate @ -80 °C overnight
(6) Transfer contents to ultracentrifuge tubes, balance with analytical balance, and spin at 20,000 xg for 30 minutes at 4 °C (see Barna lab for TLA 120.2 rotor, we have model 343778 tubes)
(7) Wash twice with 750 µL 70% EtOH, ensuring not to disrupt the pellet (don’t mix pipette). Additionally, let pellet dry for ∼10 minutes in fume hood until nearly all ethanol has evaporated
(8) Resuspend pellet in 100 µL cold HEPES buffer
(9) Quantify recovery with A260 and manufacturer’s listed molar extinction coefficient. Typical concentration is around ∼80 µM.

#### Part 2: Cell labeling

(1) Between 500K and 5M fixed cells (stored in 70%EtOH at -20C) were washed twice in 1xPBS and resuspended in 100ul of 1xPBS.
(2) 4 µL of 1 mM TCO-PEG4-TFP Ester was added to the cells, mixed by pipetting and incubated for 5 min @RT protected from light
(3) 6 µL of each methyltetrazine oligo amide (prepared in Part 1) was added, mixed by pipetting and incubated for 30 min @RT protected from light
(4) Methyltetrazine-PEG4-Amine was added for final concentration of 50 µM and Tris HCl was added to 10 mM final concentration and incubated for 5 min @RT
(5) Cells were then diluted ∼twofold with 1X PBS, spun down and washed 3x in 1xPBS
(6) Cells were resuspended in 70% EtOH in 1xPBS, and kept -20C for up to 6 months after the initial cell collection date.

### Cell plating for imaging

Cells stored in suspension at -20 C were resuspended by pipetting. Barcoded cells were combined at desired ratios in a single tube. A small volume of cells (∼5ul of cells at 10k/ul-20k/ul concentration) was spotted on a poly-lysine coated area in the center of the slide and let dry for 3-5 min. 1xPBS was then added and the slide was inspected to check for optimal cell density (ideally monolayer of cells evenly covering the entire slide). Cells were then used for IF or ORCA.

### ORCA primary probe hybridization

For ORCA DNA experiments, we used the protocol described in Mateo et al. 2019 and in detail in Mateo et al. 2020. Briefly, cells were fixed in 4% PFA in 1x PBS for 10 min. After cell plating, cells were permeabilized with 0.5% Triton-X in 1xPBS for 10min, followed by 2 washes with 1xPBS. Cells were incubated in 0.1M HCl for 5 min followed by 3 washes in 1xPBS. We then treated cells for 30min with RNAse A (10ug/ml) at 37C followed by 3 washes in 2xSSC. Cells were then incubated for 35 min in Hybridization buffer (2xSSC, 50% formamide and 0.1% Tween). Primary probes (for 2-3 Mb Hoxa and Sox2 probes, we used 15ug of probe, for 30Mb and entire chromosome tiling we used 3-4 ug of probe) diluted in hybridization solution (2xSSC, 50% formamide, 10% dextran sulfate, 0.1% Tween) were then added onto cells, covered with a coverglass, denatured for 3min at 90C and incubated overnight at 42C. The following day, cells were washed with 2xSSC, postfixed in 2%GA, 8% PFA in 1xPBS for 30min to 1h and set up on the microscope for sequential labeling and imaging. If ORCA was preceded by IF, we picked positions to match as closely as possible to cells imaged by IF. Cell barcodes were either imaged before or after probe barcodes.

RNA labeling was done as described above for DNA with the exception of the HCl, RNAse A or post fix steps.

### ORCA imaging

Samples hybridized with primary probes were imaged on the custom microscopy and microfluidics setup as previously described ^5,78^. Briefly, we used oligos complementary to the readout barcodes on primary probes carrying either a Cy5 or a 750 dye. We used strand displacement to remove an imaged barcode. We also used a Cy3 oligo to label all probes and imaged this Cy3 fiducial channel at each round of imaging. The fiducial spots were subsequently used for spot calling and registration in the image analysis pipeline.

### Immunofluorescence followed by ORCA

Plated cells were permeabilized and blocked in antibody dilution buffer (2%BSA, 0.1% Triton X - 100 in 1xPBS) for 10 min @RT. Cells were then incubated with the Geminin primary antibody (ab195047, 1:100) diluted in antibody dilution buffer for 1 h @ RT, washed 3x with 1xPBS, incubated with secondary antibody (anti-rabbit 647 Invitrogen A31573) and DAPI diluted in antibody dilution buffer for 30 min @RT, washed 3x with 1xPBS after which they were ready to image.

### Loop extrusion simulations

Loop extrusion simulations were performed using the “polychrom” package from “open2c” github project ^79^, which was developed from earlier polymer models of loop extrusion written by the Mirny lab ^21,30^, and powered by the GPU accelerated molecular simulation toolkit openMM ^80^. All simulations were run on an NVIDIA Titan Xp card. polychrom uses a Langevin approach to simulate the dynamics of a polymer under several user defined energy constraints. Complete details of the energetic constraints and other parameters are specified in the python simulation scripts included in our github repository for this project: https://github.com/BoettigerLab/Hafner2022/, and described in brief below. These can be run independently using polychrom, which may be downloaded from the polychrom github page: https://https://github.com/open2c/polychrom.

We simulated the chromosomes as 6 polymers confined in a spherical geometry representing the nucleus. Each polymer was 5,000 monomers long, corresponding to 30 kb/monomer for 150 Mb chromosomes, on par with the probe size used at our 3 Mb, 30 Mb, and whole-chromosome (150 Mb) experiments. Each simulation was run in 600 independent replicates, for sufficient time that the final 3D structures were uncorrelated with the starting structure at all length scales and that the pattern of TADs and loops could be reproduced from averaging single replicates over time identically as to averaging over replicates (see below for loop extrusion and TADs). For high resolution experiments, we conducted additional simulations, consisting of 2 polymers constrained in a spherical volume, each 50,000 monomers long (3 kb/monomer). These simulations ran substantially more slowly, and even after running 10 fold longer the structure at the very longest length scales was not completely uncorrelated from the initialization conditions, though the structures at shorter length scales had reached a stationary distribution. For these reasons, we used these high resolution simulations only to model the submegabase and not whole chromosome organization.

Loop extrusion was simulated using the previously described sequential 1D and 3D simulations, implemented by *polychrom.* Briefly, the 1D simulations simulate the loading, unloading, and motion of loop extruders with left and right arms that walk bi-directionally on chromatin and interactions with extrusion blockers. These are followed by 3D simulations, in which short harmonic bonds between the left and right arms of the extrusion factors impose the loop. Loop extrusion blocking sites, simulating CTCF, were distributed in the pattern shown in Fig 6D. This 3 Mb pattern was replicated back to back to produce the 150 Mb chromosome pattern. Cohesins were loaded with an average spacing of 300 kb, and a lifetime of corresponding to a maximum extrusion length of 600 kb if not stalled. The confinement radius for the nucleus was chosen so 20% of the nuclear volume was filled with polymer. Confinement volumes 0.02% were used to represent the unconstrained conditioned and 0.2 the 10% of relaxation volume condition. A hard-sphere repulsion energy of 6 was used in all simulations except simulating a higher degree of chain crossing, which used a value of 3. No additional attractive or repulsive interactions between monomers was used in the simulations, which do not attempt to recapitulate the A/B compartmentalization. See code available on github link above for details.

It should be emphasized that these are coarse-grained simulations, intended to capture the qualitative, emergent properties of a small number of physical processes – such as loop extruders moving on a large polymer in a confined volume. The utility of these coarse grained physics models is not to capture as many features of reality as possible in a common model, but rather to reproduce key patterns in the experimental data from a minimal mechanistic hypothesis ^81,82^. By using as few features from reality as possible in the model rather than as many as possible, we can develop deeper intuition about the underlying processes. As a consequence of coarse-graining abstraction, the precise relation between simulation units, monomers to kb, cohesin spacing to kb, monomer radii to nanometer, etc, are approximate. We did not undertake a comprehensive sweep of the parameter space of the model. The intensive computational time for the thousands of independent simulations limit such analysis, and showing that the multiscale qualitative effects of cohesin chromatin folding can be captured by loop extrusion did not require it.

### Image processing and quantification

#### Image processing and spot calling for ORCA data

Image processing (drift correction and localization of spots) analysis was performed as described in ^5,78^.

#### Cell segmentation

We used the CellPose ^83^ for segmenting individual nuclei.

#### Geminin IF and cell size quantification

For Geminin IF quantification and the quantification of nuclear size, we performed all calculations on the experiments for the Hoxa and Sox2 domains for 2.2 and 2.8 Mb respectively where cells were stained for Dapi and Geminin prior to the ORCA experiments. Nuclear segmentation was done on Dapi staining using CellPose. The resulting cell masks were then used to quantify levels of Geminin immunofluorescence by computing the mean intensity per nucleus. The Geminin low population was defined as the lowest 25% of the geminin expression level across all stained cells within the experiment (combined the E14 parental cell line together with CTCF-AID and Rad21-AID auxin treated and untreated cells). For nuclear area quantification, we used the total number of pixels in each segmented mask in this experiment.

#### Demultiplexing cell barcodes

For each fiducial spot, we quantified the intensity of each cell barcode signal. To normalize for differences in brightness across cell barcodes, we then balanced the intensities across all values. We used the nearest neighbors approach to classify each spot into groups based on the values for all cell barcodes for that spot.

#### RNA FISH data analysis

RNA FISH data was quantified similarly to ^5^. Briefly, images were maximum-z-projected and flatfield corrected. Foci corresponding to single mRNAs were identified by using a local-maxima search with manually defined thresholds. Foci positions were then overlaid with cell segmentation masks from CellPose to compute single-cell transcript mRNA counts.

### ORCA analyses

#### Merge and filtering of ORCA data

For the ∼3 Mb scale data, we had 2 technical replicates (performed on the same batch of collected and barcoded cells) and a biological replicate (independently thawed, treated, fixed and barcoded cells). Replicates showed a high degree of reproducibility and showed identical patterns of TADs and loops and were comparable in median distance (Supplement 1). We thus merged the data for all subsequent analyses. Barcodes with low hybridization efficiency (< 10 %) were filtered on the merged data. An additional filter was done on a dataset basis to remove obvious artifacts such as failed strand displacement events.

#### Insulation score

Insulation score was calculated on contact frequency maps. We used a window size of 8 barcodes (8 * 30 kb steps size = 240 kb). For each position along the diagonal, we calculated the sum of contacts in the upstream and downstream window. To be able to compare values across conditions we then divided the values by the mean insulation score in the untreated condition.

#### Three-way contact analyses

For the three-way CTCF contact analyses in Fig 2, we picked the 3 CTCF sites for *Hoxa* (barcodes 20, 34, 54) and *Sox2 regions* (barcodes 42, 46, 70). We calculated the frequency of chromosomes where the distance between sites 1-2 and 2-3 were both below 200 nm. This frequency was normalized for detection efficiency for these pairs. We then shifted barcode positions 8 steps (with each step= 30 kb) in each direction and performed the same calculation for each shift. The expected frequencies were generated by multiplying the frequency for the pairs (frequency(1,2) * frequency (2-3)) and dividing by their detection efficiency.

We performed bootstrapping to estimate the confidence intervals for the contact frequencies. Bootstrapping was done by randomly sampling with replacement from the data. The height bars in the barplot (Fig. 2, Supplement 3) correspond to the mean and the error bars correspond to the 25th and 75th quantiles. Thus, errorbars that do not overlap reflect contact-frequencies that were distinct in 94% of resamplings (94 = 1-0.25^2^). We used 200 resampling draws in our bootstrapping, and noted more resamplings made little difference on the spread.

#### Compartment analysis using Pearson’s Matrix

Compartmentalization was analyzed in the ORCA data starting from the median pairwise distance matrices, **O**, for both the 30 Mb scale and whole chromosome (150 Mb) data. The “expected” distance was computed by averaging the 3D distance for all loci pairs which had the same genomic spacing. This produces a matrix, **E**, with a smooth decay from the main diagonal. We then divided the observed distances by the expected distances to create a new matrix, **N**, N*_i,j_* = O*_i,j_*/E*_i_*,. Finally, we computed the Pearson Matrix **P**, such that P*_i,j_* is Pearson’s correlation coefficient between row *i* and column *j* of matrix **N**. This parallels the original analysis of compartmentalization described for Hi-C, though using our distance measures in place of the contact-frequency per bin ^38^.

We note that the Hi-C and ORCA compartment analysis were computed with an important difference. The Hi-C compartment analysis at the 30 and 150 Mb scale uses reads from every part of the 30Mb (150 Mb) interval. In contrast, the ORCA data looks exclusively at particular 30 kb elements that skip through that domain at 1 Mb or 5 Mb intervals. Thus, substantial parts of the genome driving the correlation signal in the Hi-C data are not included. We note that our calculation using Hi-C data that focused only on the interactions between the 30 kb windows covered by the ORCA experiments were too sparsely populated to permit detection of compartments. Expanding these to 120 kb bins resulted in only minor improvement, and thus we elect to show the Hi-C compartmentalization computed using all the data. Thus, some qualitative differences in the position/distribution of the compartment boundaries likely reflects the contributions of genomic sequences not covered in our imaging experiments, though the sparsity of the Hi-C data makes this difficult to quantify precisely.

### Published genomic data

#### Hi-C data

We examined Hi-C experiments from ^16,19^ for CTCF degradation, from ^18^ for Rad21 degradation, from ^84^ for untreated cells as these were all generated in mESCs. For Nora et. al and Rhodes et al. datasets, we loaded the processed Hi-C .cool files from either GEO (GSE98671) or ArrayExpress (E-MTAB-7816) respectively. We used Cooler ^85^ to export normalized tables directly using the cooler ‘dump’ command for the coordinates that match our probes. For the Kubo et al. dataset, we loaded the processed .hic files from GEO (GSE94452), uploaded the .hic files to Juicebox ^86^ and exported the normalized tables. For Bonev et al. dataset, we exported the normalized tables for the regions of interest directly from Juicebox. We then downsampled the data by averaging bins to match the resolution of the probes used for ORCA (30kb for the ∼2.5Mb scale, 1Mb for the ∼30Mb scale and 5Mb for the ∼150Mb scale probes).

#### ChIP data

We used CTCF and Ring1b ChIP-seq data from ^84^ downloaded from GEO in the bigwig format (GSE96107_ES_Ring1B.bw and GSE96107_ES_CTCF.bw), converted to bedGraph format using bigWigToBedGraph program. We then mapped the genomic coordinates with coordinates for the Hoxa and Sox2 probes. For the CTCF density plot, we averaged the signal for each barcode (spanning 30kb regions).

## Acknowledgements

We thank Max Imakaev and Neil Chowdhury for code sharing and feedback on the polymer simulations; the 4DN community for support and the opportunities to present this work; Andrea Cosolo (4DN) for help with formatting and uploading the data to the 4DN server; S. Murphy for support and comments on the text and Figures; current and former members of the Boettiger lab for comments and discussion. This work was supported by NIH grant U01DK127419, the Packard Foundation, and a Beckman Young Investigator’s Award to A.N.B. M.P. was supported by a postdoctoral fellowship from the Damon Runyon Cancer Research Foundation. A.H. was supported by Walter and Idun V. Berry Postdoctoral fellowship. S.E.B. was supported by the Stanford Graduate Fellowship. A.H. and A.N.B. designed the experiments. A.H., M.P. and S.E.B. performed the experiments. E.N. contributed key reagents. A.H. and A.N.B. performed analysis and interpreted the results. A.H. and A.N.B., wrote the manuscript with input from E.N. S.E.B. and M.P.

## Data and code availability

All probe coordinates and data tables are in the process of being annotated and formatted for upload on the 4DN public access server. Probe libraries used in this work can be viewed through the UCSC genome browser: https://genome.ucsc.edu/s/toniai/mm10_Hafner2022

Code to reproduce the simulations described in this work is available at https://github.com/BoettigerLab/Hafner2022/. These simulations require the polychrom simulation tools, available at: https://https://github.com/open2c/polychrom. Additional information on the simulations is included in the **Methods**.

## Figures and figure legends

**Extended Data Fig.1.**
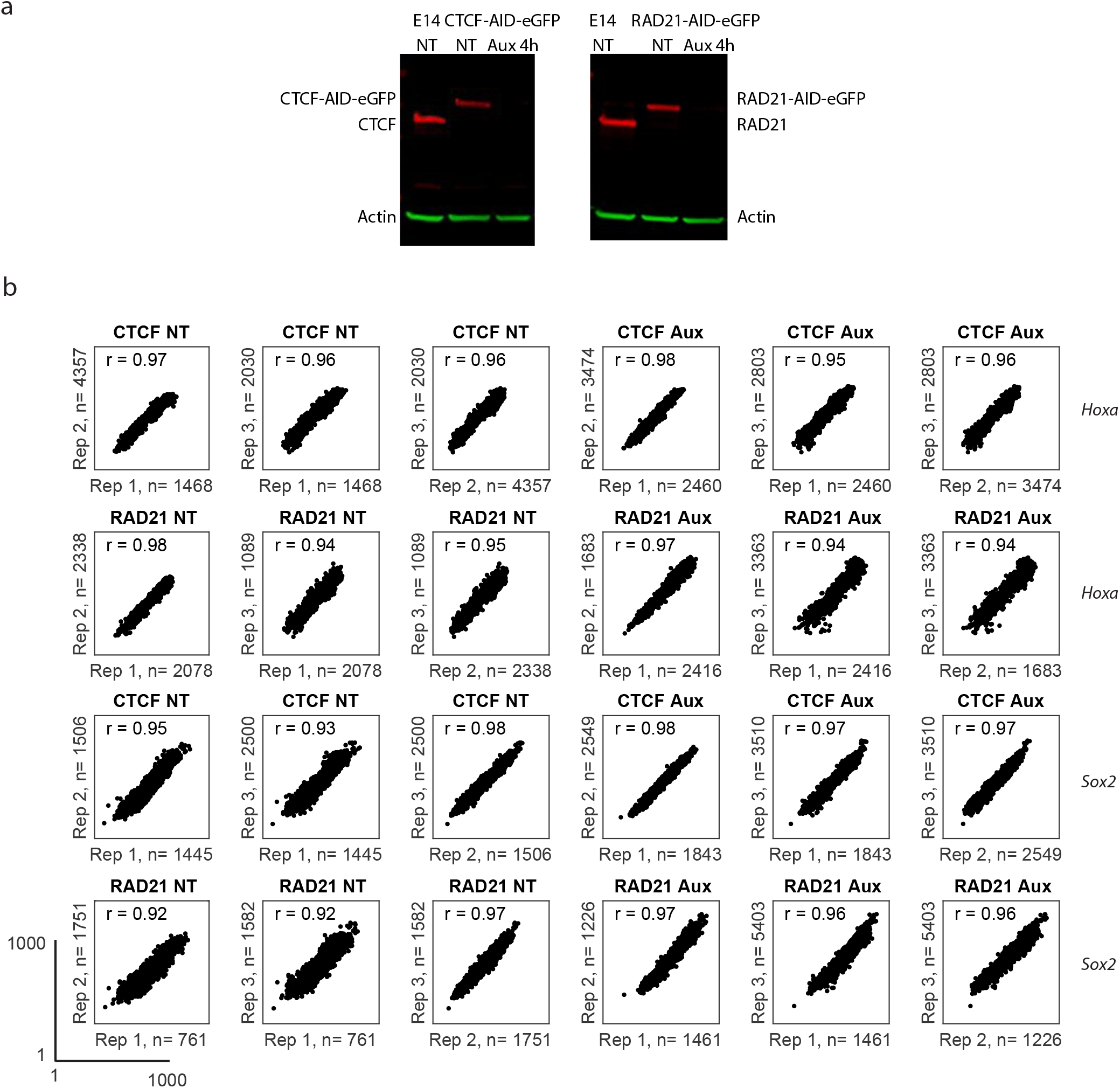
CTCF and cohesin can be depleted after auxin addition and reproducibility of ORCA replicates (related to Figure 1) **a.** Western blot labeling Actin (loading control) and either CTCF or Cohesin in the parental E14 cell line and in the CTCF-AID-eGFP or RAD21-AID-eGFP cells, with or without auxin treatment. **b**. Scatter plots of inter-barcode distances, in nanometers, comparing reproducibility among three replicates labeling the *Hoxa* or *Sox2* domains. Median distance is plotted for each barcode pair against the pairwise median distance in a different replicate. Pearson’s correlation coefficient, r, is shown on the plot. The lower left plot shows the axis scale and the same scale was used for all plots.

**Extended Data Fig. 2.**
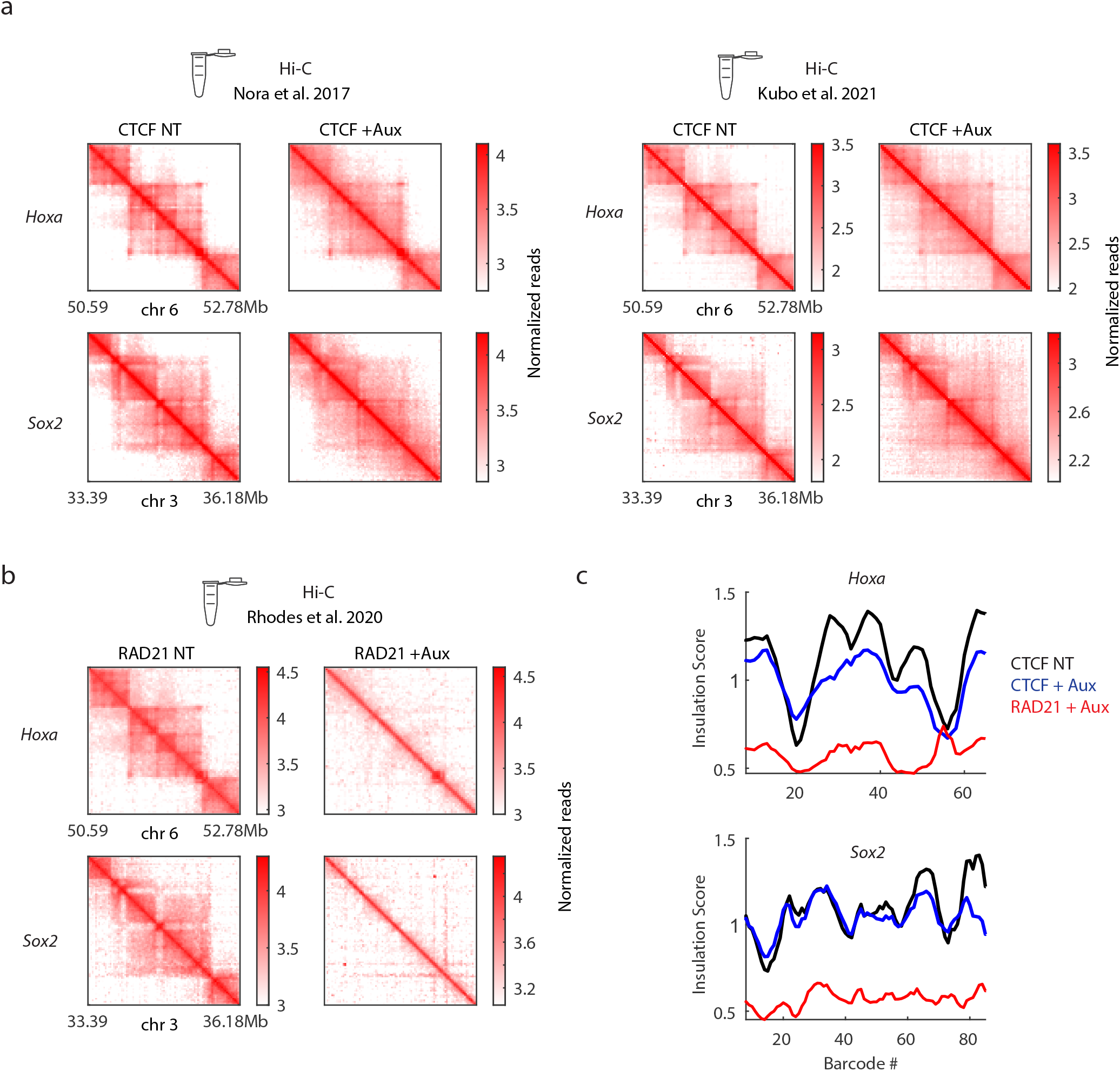
Published Hi-C data for the Hoxa and Sox2 domains qualitatively agrees with ORCA (related to Figure 1) **a-b**. Hi-C data^16,18,19^ for the *Hoxa* and *Sox2* domains corresponding to regions imaged with ORCA. **c.** Insulation score calculation for *Hoxa* and *Sox2* domains done on contact frequency maps of CTCF untreated, CTCF auxin-treated and RAD21 auxin-treated conditions.

**Extended Data Fig. 3.**
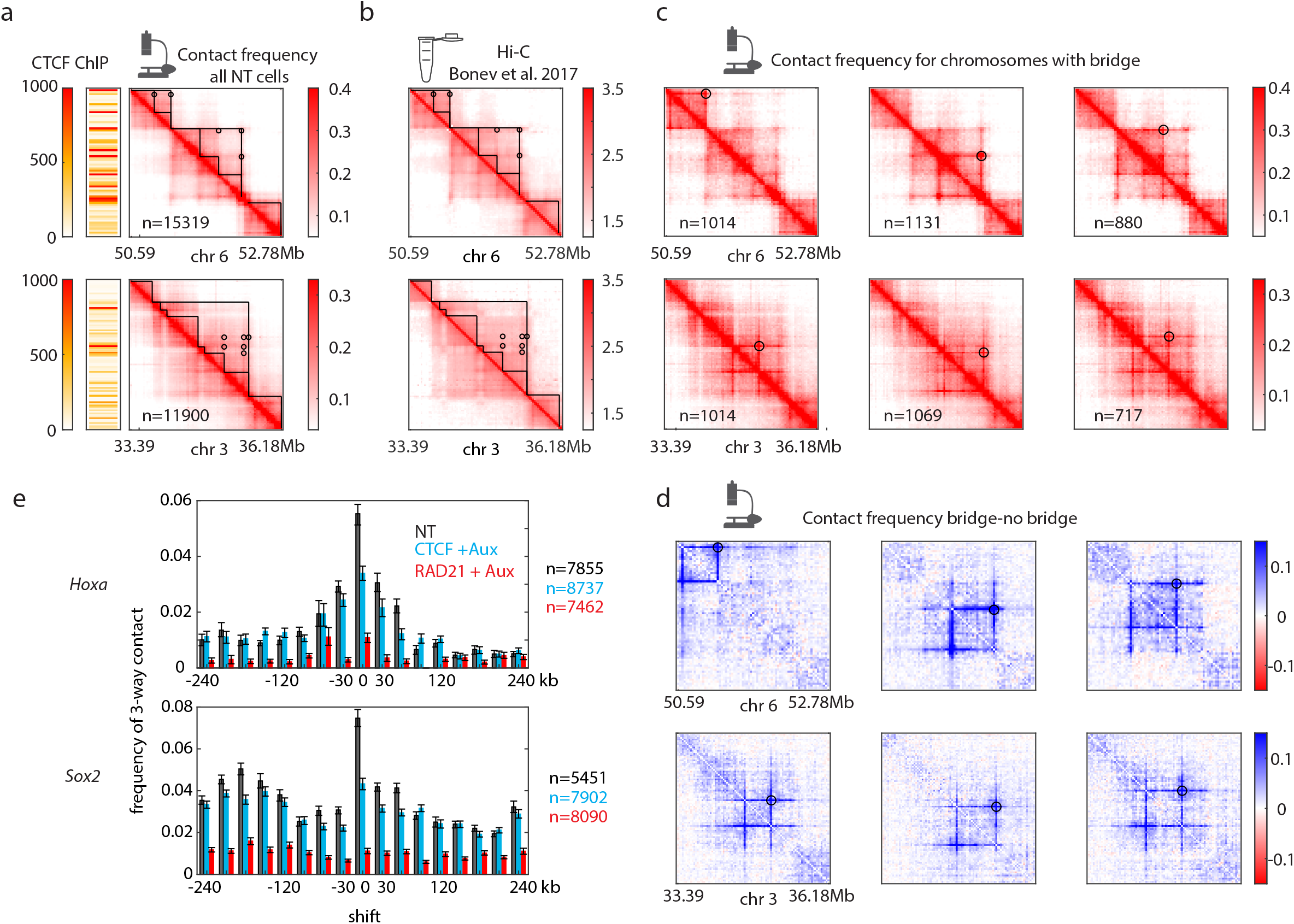
“Bridges” in ORCA and Hi-C (related to Figure 2). **a.** Contact frequency maps from untreated cells. Circles denote visible loops that bridge across TAD boundaries. The heatmap on the left of the contact frequency maps shows the density of CTCF ChIP-seq signal. **b**. Hi-C contact frequency for untreated mESCs from Bonev et al. 2017 ^84^ showing the same bridging contacts as in (a). **c**. Contact frequency map for cells with different bridging contacts. **d.** Difference in contact frequency maps between cells that have the bridging contacts and cells that do not. **e**. Frequency of the 3-way contact between the positions corresponding to the CTCF-marked boundaries relative to positions not near a boundary, as in Fig. 2f.

**Extended Data Fig. 4.**
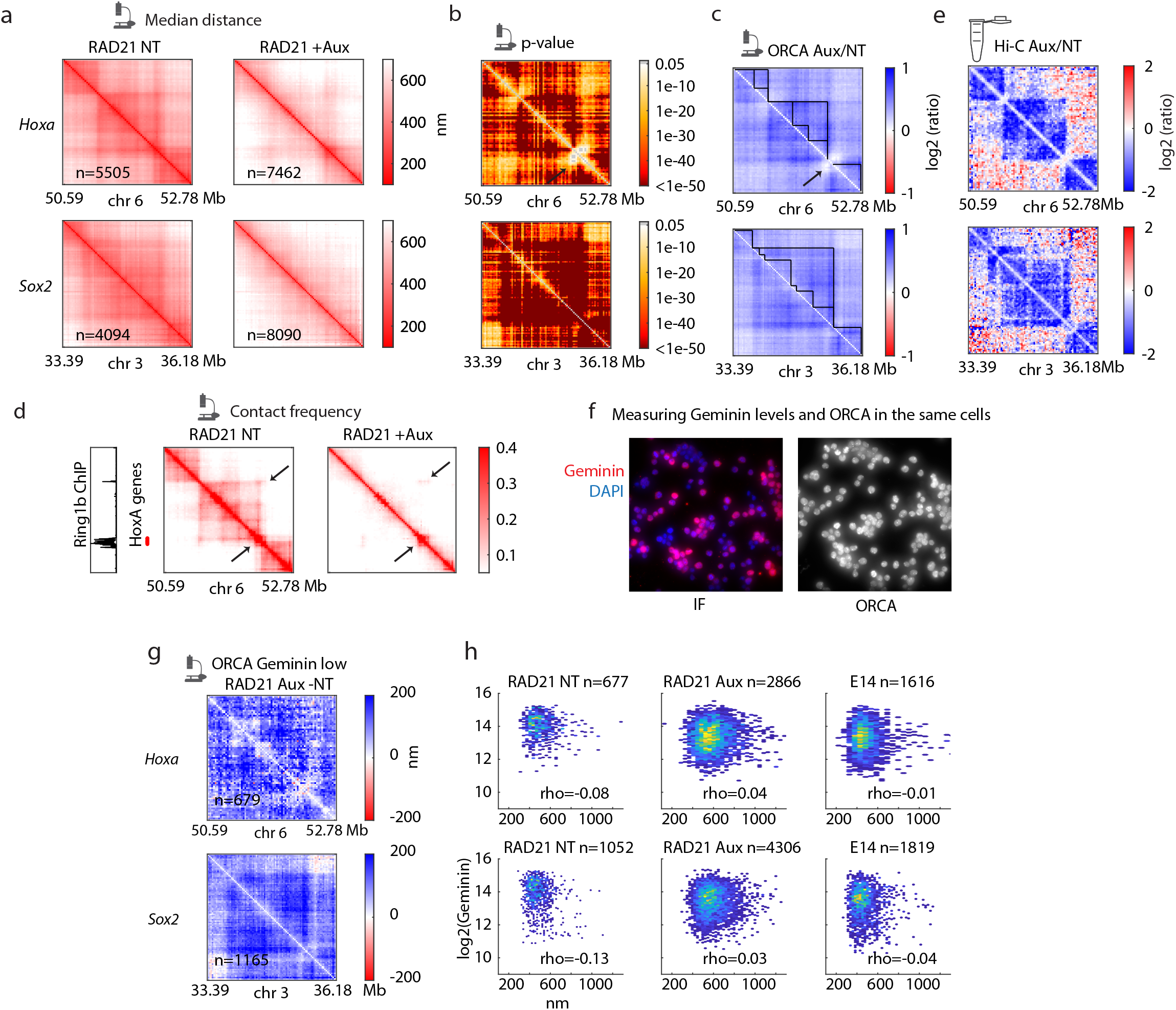
Cohesin depletion leads to expansion at the ∼3 Mb scale (related to Figure 3). **a.** Median distance maps for the *Hoxa* and *Sox2* domains in untreated or auxin treated cells. **b**. Maps of P-values computed using ranksum for each pairwise distance between auxin treated and untreated cells. Arrow indicates the only regions other than the diagonal that did not significantly change upon auxin treatment. **c**. Log2 ratio of median distance between auxin-treated and untreated cells. Black lines indicate TAD outlines and the black arrow indicates the region that did not change upon auxin treatment. **d**. Contact fraction maps for the *Hoxa* region and the corresponding Ring1B (member of the polycomb complex) ChIP signal from ^84^ as well as the position of *Hoxa* genes. Arrows point to Ring1B coated regions on the heatmaps. **e.** Log2 ratio of Hi-C data between RAD21 auxin-treated and untreated cells shows a decrease in intra-TAD contacts and gain of contacts between TADs. **f.** Example field of view image of DAPI and immunofluorescent staining of Geminin and ORCA fiducial staining for the same cells. **g**. Difference in median distance between the Geminin-low population for the RAD21 auxin treated cells and all RAD21 untreated cells. **h**. Scatter plot of median distance (over all pairs) in each chromosome relative to Geminin levels in the same cell. Spearman’s correlation coefficient is indicated for each condition.

**Extended Data Fig. 5.**
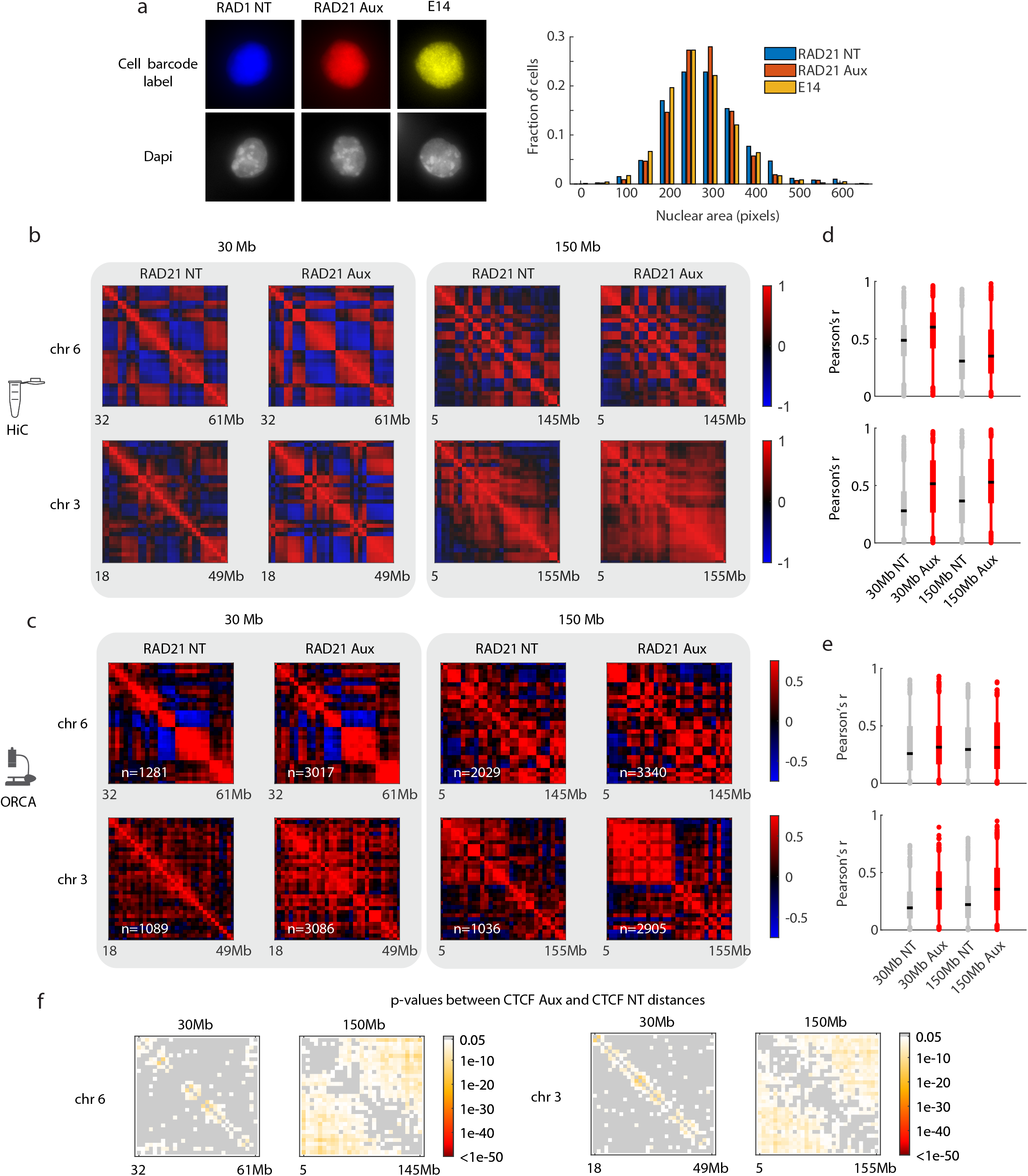
Quantification of nuclear size and compartments after depletion of cohesin (related to Figure 4) **a.** Example images of individual cells labeled with DAPI and the cell surface marker for the parental E14 cell line, RAD21 NT or RAD21 Aux cells and the quantification of cell sizes across all cells (n=1729 cells for RAD21 NT, n=7172 cells for RAD21 +Aux, n=3435 for E14). **b**. Hi-C ^84^ Pearson correlation maps for the 30 Mb and 150 Mb regions corresponding to regions imaged by ORCA and the distribution of Pearson correlation coefficients. **c**. ORCA Pearson correlation maps for the 30 Mb and 150 Mb probes in RAD21 untreated or auxin treated cells. **d.** Distribution of Pearson correlation coefficients for Hi-C data. **e**. Distribution of Pearson correlation coefficients for ORCA data. **f**. P-values computed using ranksum for each pair between auxin-treated and untreated cells.

**Extended Data Fig. 6.**
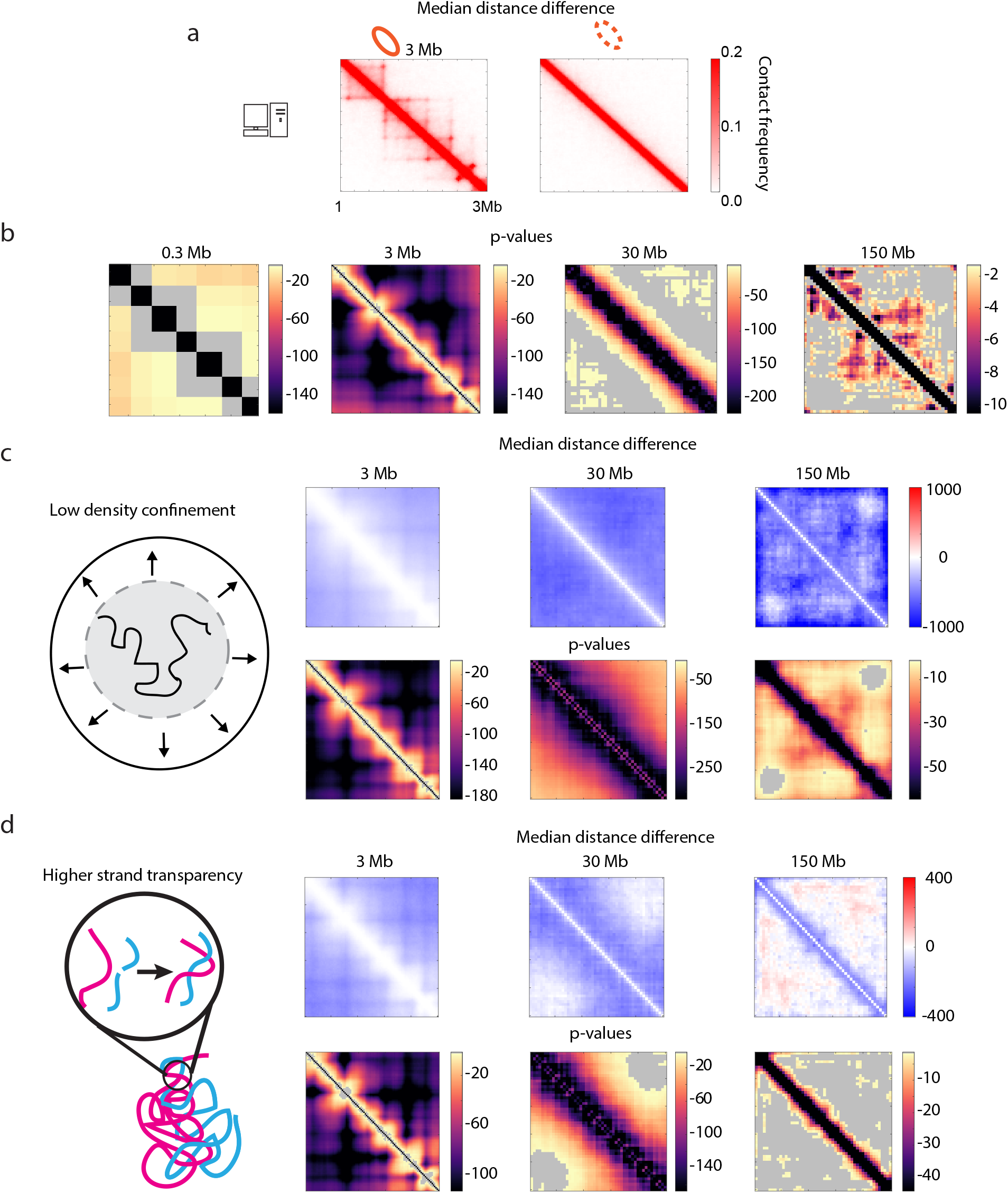
Polymer simulations with lower nuclear confinement and increased strand crossing (related to Figure 5) **a.** Contact frequency of simulated polymers with or without loop extrusion factors. **b**. p-values comparing simulated polymers with or without loop extrusion factors at different genomic scales. **c-d**. Median distance difference maps and p-values for simulations with or without loop extrusion factors at different scales for simulations with low density confinement (c) or with more strand crossing (d).

**Extended Data Fig. 7.**
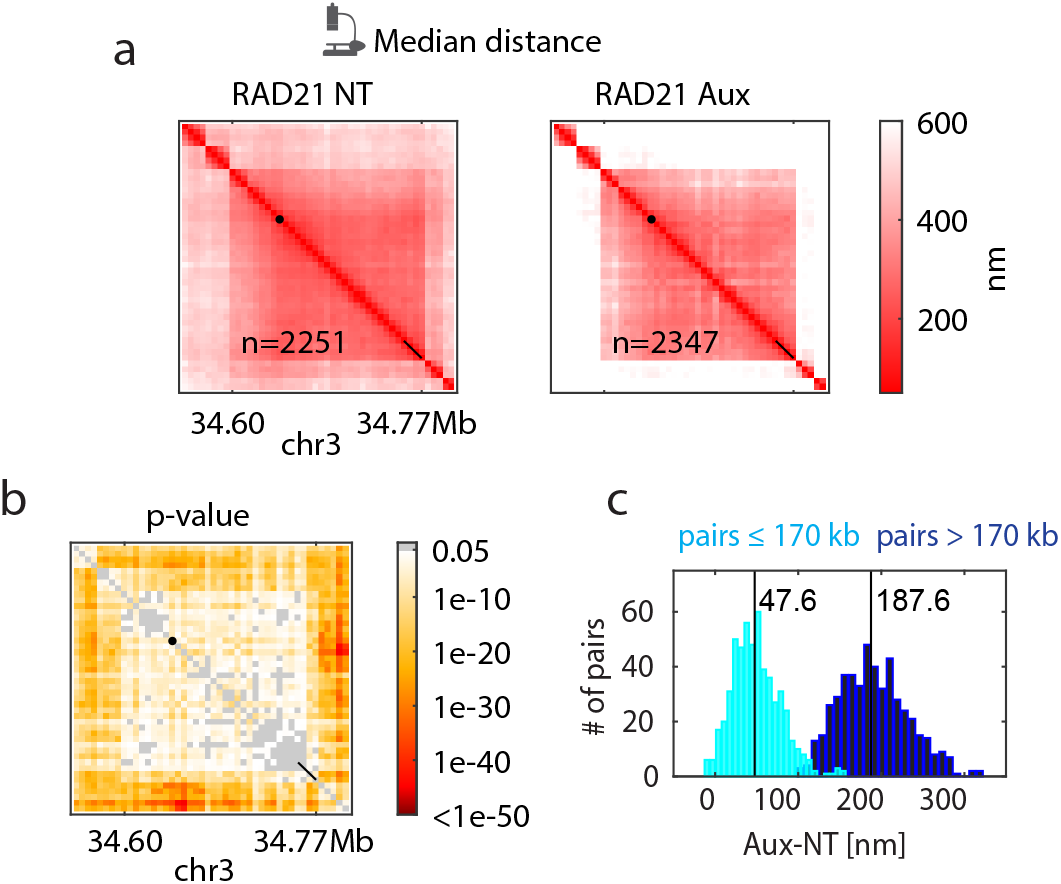
Additional ORCA data for the fine-scale Sox2 region (related to Figure 5) **a.** Median distance and **b**. p-values for the *Sox2* 170 kb domain with genomically distant regions imaged by ORCA. **c**.Histogram of pairwise distance differences in median distance. In cyan are pairs that are <170 kb in genomic distance relative to >170 kb in dark blue.

**Extended Data Fig. 8.**
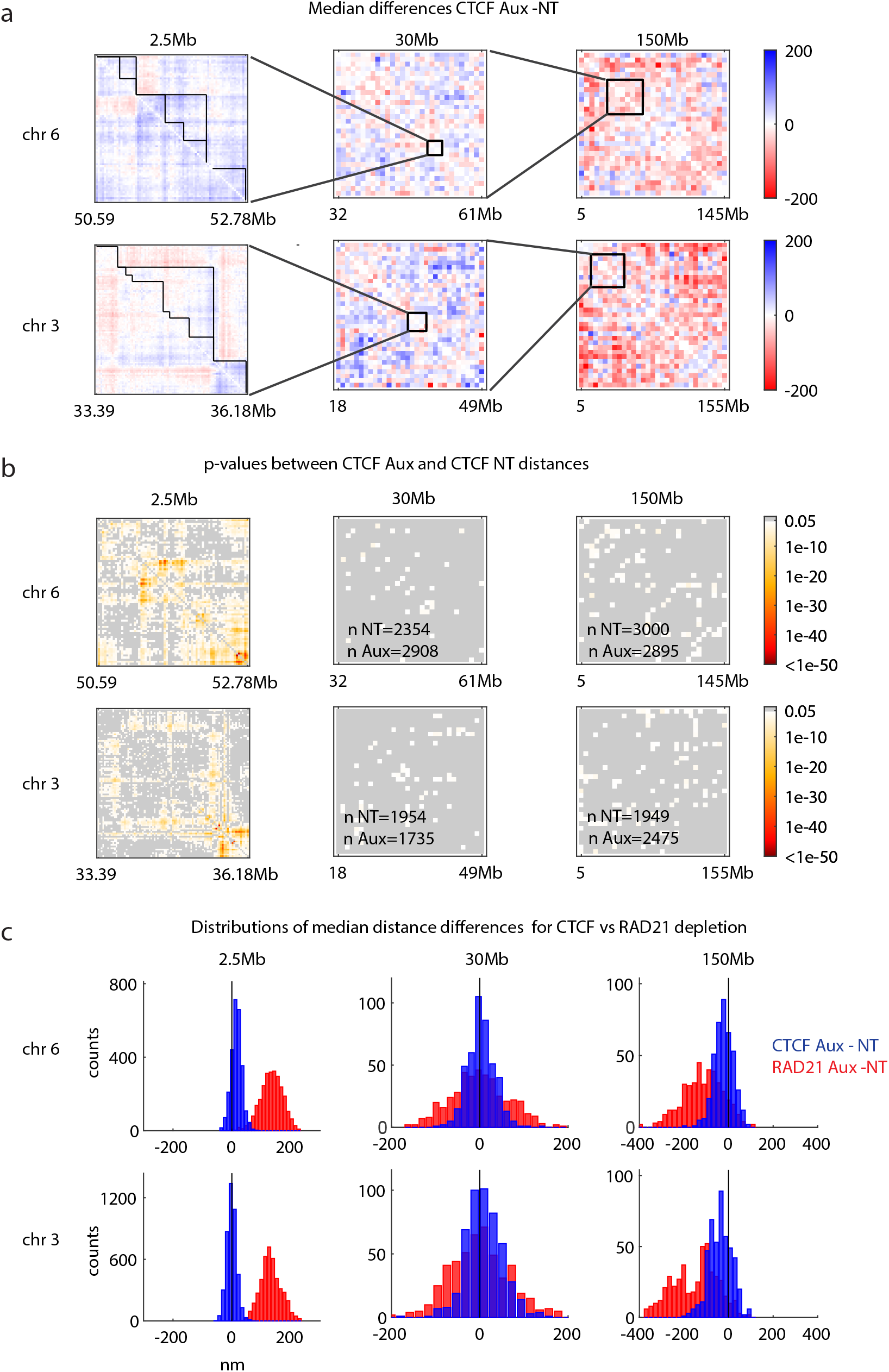
CTCF depletion leads to less change in physical distance than cohesin depletion (related to Figure 6) **a.** Median distance difference maps for CTCF Aux - CTCF NT cells at different scales of imaging. **b**. P-values computed using ranksum for each pair between CTCF auxin-treated and untreated cells at different genomic scales. **c**.Comparison of median distance differences between auxin-treated and untreated cells for either CTCF (blue) or RAD21 (red).

